# Myeloid cell-driven nonregenerative pulmonary scarring is conserved in multiple nonhuman primate species regardless of SARS-CoV-2 infection modality

**DOI:** 10.1101/2021.11.28.470250

**Authors:** Alyssa C Fears, Brandon J Beddingfield, Nicole R Chirichella, Nadia Slisarenko, Stephanie Z Killeen, Rachel K Redmann, Kelly Goff, Skye Spencer, Breanna Picou, Nadia Golden, Duane J Bush, Luis M Branco, Matthew L Boisen, Hongmei Gao, David C Montefiori, Robert V Blair, Lara A Doyle-Meyers, Kasi Russel-Lodrigue, Nicholas J Maness, Chad J Roy

## Abstract

The novel coronavirus SARS-CoV-2 has caused a worldwide pandemic resulting in widespread efforts in development of animal models that recapitulate human disease for evaluation of medical countermeasures, and to dissect COVID-19 immunopathogenesis. We tested whether route of experimental infection substantially changes COVID-19 disease characteristics in two species (*Macaca mulatta*; rhesus macaques; RM, *Chlorocebus atheiops*; African green monkeys; AGM) of nonhuman primates. Species-specific cohorts of RM and AGM Rhesus macaques (*Macaca mulatta*, RMs) and African green monkeys (*Chlorocebus aethiops,* AGMs) were experimentally infected with homologous SARS-CoV-2 by either direct mucosal instillation or small particle aerosol in route-discrete subcohorts. Both species demonstrated equivalent infection initially by either exposure route although the magnitude and duration of viral loading was greater in AGMs than that of the RM. Clinical onset was nearly immediate (+1dpi) in mucosally-exposed cohorts whereas aerosol-infected animals began to show signs +7dpi. Myeloid cell responses indicative of the development of pulmonary scarring and extended lack of regenerative capacity in the pulmonary compartment was a conserved pathologic response in both species by either exposure modality. This pathological commonality may be useful in future anti-fibrosis therapeutic evaluations and expands our understanding of how SARS-CoV-2 infection leads to ARDS and functional lung damage.

## Introduction

SARS-CoV-2, the viral pathogen responsible for the current worldwide pandemic, has caused over 737,000 deaths in the United States and over 4.9 million deaths worldwide^1, 2^. Pathogenesis of this disease in humans includes pneumonia accompanied by shortness of breath, with a subset of affected individuals experiencing acute respiratory distress syndrome (ARDS) and death^3^. Symptoms have been reported to continue beyond resolution of viral persistence, including fatigue, dyspnea, anosmia and headache^4^. Symptoms lasting longer than four weeks past this point being termed post-acute COVID-19 syndrome (PACS), or ‘long COVID’^5^.

The exploratory development of COVID-19 disease models using various animal species continues to be goal to combat this pandemic and investigations have led to highly useful test systems for evaluation and pathogenesis studies^6^. Accordingly, multiple nonhuman primate (NHP) species have been experimentally infected with SARS-CoV-2, with most studies focusing on *Chlorocebus aethiops* (African green monkeys, AGMs) or *Macaca mulatta* (Rhesus macaques, RMs). Other examples of species studied include *Macaca nemestrina* (Southern pigtail macaque)^7^, *Macaca leonina* (Northern pigtail macaque)^8^, *Macaca fascicularis* (Cynomolgus macaque)^9^, *Callithrix jacchus* (Common marmoset) and *Papio hamadryas* (Baboon)^10^ . Most of these models involve installation of the virus directly to mucosal surfaces^11, 12^, though some have included the aerosol modality of exposure^13^. Overwhelmingly, the NHP model of SARS-CoV-2 infection results in a mild to moderate disease, with only one study reporting euthanasia criteria being met post challenge^14^. The RM model of disease, utilized for vaccination^15, 16^, re-challenge^17^, and therapeutic^18^ studies, results in disease resolving within three weeks post challenge^9^, though some evidence of longer term viral replication has also been reported^19^. AGMs have been utilized for many similar respiratory-based viral diseases including SARS-CoV-1^20^, parainfluenza virus^21^ and Nipah virus^22^. Their use as SARS-CoV-2 infection models has resulted in observed mild respiratory disease like RMs, but with prolonged shedding of viral RNA^11, 13^.

The most common severe disease outcome in humans infected with SARS-CoV-2 is ARDS. Fibroproliferative disease following resolution of ARDS is the most common cause of death in patients, with up to 61% of autopsies showing signs of pulmonary fibrosis and 25% of ARDS survivors show evidence of restrictive lung disease with long lasting morbidity^23–25^. Within the COVID patient cohort, 42% who develop severe pneumonia will progress to ARDS, with fatal cases generally presenting with pulmonary fibrosis at autopsy^26, 27^. ARDS in COVID is characterized by a myeloid cell migration into the lung ^28^, with early NHP studies showing infiltration of CCR2+ myeloid cells.

These studies did not allude to the mechanism by which the cellular kinetics or phenotypes specifically correlated with the pathologic sequalae of pulmonary scarring from this viral disease^29^.

In this study, we tested whether NHP species or exposure modality functionally changes disease course and progression. RM and AGM cohorts are compared experimentally infecting either species via direct mucosal route (intratracheal/intranasal) or small particle aerosol modality to a low passage SARS-CoV-2 archival (WA1/2020) strain.

We demonstrate that infection by small particle aerosol results in slower development of signs of disease and immune response to viral infection in both species. The AGMs revealed longer viral dynamics in the respiratory compartment and longer-term elimination in the gastrointestinal system when in contrast to the RM. Myeloid cell kinetics and phenotypes were defined among the entire cohort to investigate whether variables associated with experimental infection or differences in species susceptibility attribute to the severe outcome of SARS-CoV-2 infection. Myeloid cell infiltration and anti-inflammatory phenotype correlating with decreased pulmonary scarring in either species. Lack of regenerative activity in the lung was present in both species post resolution of most of the viral RNA, indicating the NHP model of SARS-CoV-2 infection can be utilized during anti-fibrosis therapeutic development and evaluation and has potential utility in evaluation of post-acute COVID sequelae.

## Methods

### Animal cohort and procedures

A total of 16 NHPs were utilized for this study, between 4 and 11 years old, with most being 7 years of age. All RMs used in this study were captive bred at TNPRC. Four individuals of each species were challenged with SARS-CoV-2 USA_WA1/2020 (World Reference Center for Emerging Viruses and Arboviruses, Galveston, TX), by small particle aerosol, with a mean delivered dose of 1.5 x 10^4^ TCID_50_. The animals exposed to SARS-CoV-2 by aerosol were individually exposed using well-established methodologies as reported in the literature^14, 30^. The other four animals of each species were challenged via a combination of intratracheal and intranasal administration (IT/IN), herein referred to as ‘multiroute’, with a delivered dose of 2.0 x 10^6^ TCID_50_ (Table S1). Pre- and post-exposure samples were taken from blood, mucosal (pharyngeal, nasal, rectal) swabs, and BAL supernatant. Chest X-rays were also performed regularly during the study (Figure 1). Animals were monitored for signs of disease throughout the study, with no animals reaching euthanasia criteria. At necropsy, mucosal samples were taken, as well as tissues placed in Trizol or z-fix or fresh frozen for later examination.

**Figure 1:**
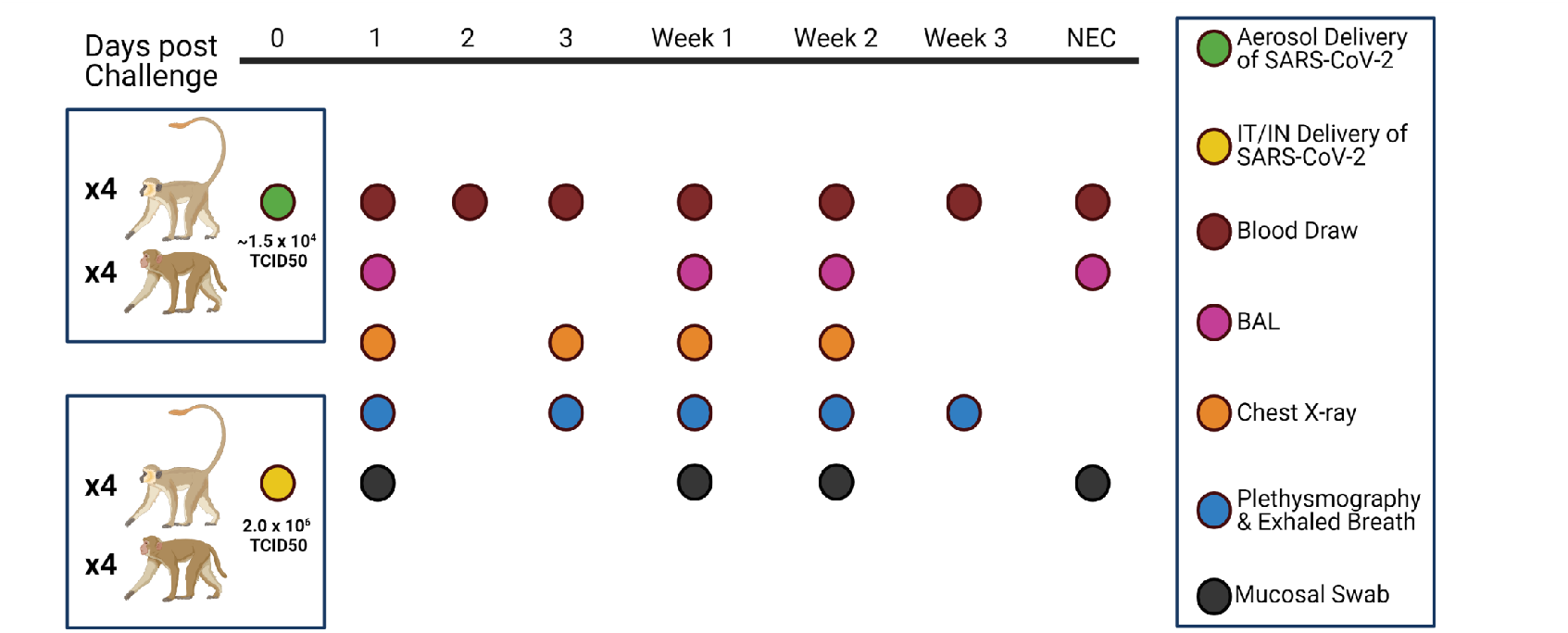
Study Design. SARS-CoV-2 was delivered via aerosol or IT/IN (multiroute) to RMs or AGMs at the noted doses. Animals were followed for 4 weeks, with biosampling performed as indicated above. Figure created in Biorender.com.

### Ethical Statement

The Tulane University Institutional Animal Care and Use Committee approved all procedures used during this study. The Tulane National Primate Research Center (TNPRC) is accredited by the Association for the Assessment and Accreditation of Laboratory Animal Care (AAALAC no. 000594). The U.S. National Institutes of Health (NIH) Office of Laboratory Animal Welfare number for TNPRC is A3071-01. Tulane University Institutional Biosafety Committee approved all procedures for work in, and removal of samples from, Biosafety Level 3 laboratories.

### Isolation of Viral RNA

RNA was isolated from non-tissue samples using a Zymo Quick RNA Viral Kit (#R1035, Zymo, USA) or Zymo Quick RNA Viral Kit (#D7003, Zymo, USA) for BAL cells, per manufacturer’s instructions. RNA was eluted in RNAse free water. During isolation, the swab was placed into the spin column to elute the entire contents of the swab in each extraction. BAL supernatant was extracted using 100 μL. Viral RNA from tissues was extracted using a RNeasy Mini Kit (#74106, Qiagen, Germany) after homogenization in Trizol and phase separation with chloroform.

### Quantification of Viral RNA using Quantitative Reverse Transcriptase PCR

Isolated RNA was analyzed in a QuantStudio 6 (Thermo Scientific, USA) using TaqPath master mix (Thermo Scientific, USA) and appropriate primers/probes (Table S2) with the following program: 25°C for 2 minutes, 50°C for 15 minutes, 95°C for 2 minutes followed by 40 cycles of 95°C for 3 seconds and 60°C for 30 seconds. Signals were compared to a standard curve generated using *in vitro* transcribed RNA of each sequence diluted from 10^8^ down to 10 copies. Positive controls consisted of SARS-CoV-2 infected VeroE6 cell lysate. Viral copies per swab were calculated by multiplying mean copies per well by amount in the total swab extract, while viral copies in tissue were calculated per μg of RNA extracted from each tissue.

### Live Virus Quantification

Median Tissue Culture Infectious Dose (TCID_50_) was used to quantify replication-competent virus in swabs and BAL supernatant. VeroE6 ells were plated in 48-well tissue culture treated plates to be subconfluent at time of assay. Cells were washed with serum free DMEM and virus from 50 μL of sample was allowed to adsorb onto the cells for 1 hour at 37°C and 5% CO_2_. After adsorption, cells were overlayed with DMEM containing 2% FBS and 1% Anti/Anti (#15240062, Thermo Scientific, USA). Plates were incubated for 7-10 days before being observed for cytopathic effect (CPE). Any CPE observed relative to control wells was considered positive and used to calculate TCID_50_ by the Reed and Muench method ^31^.

### Detection of Antibodies in Serum

The ability of antibodies in serum to disrupt the binding of the receptor binding domain (RBD) of SARS-CoV-2 spike protein to Angiotensin Converting Enzyme (ACE2) was assessed via the Surrogate Virus Neutralization Test (GenScript# L00847) using the included kit protocol modified per the following: Serum samples were diluted from 1:10 to 1:21,870 to determine an IC_50_ for RBD/ACE2 binding. Pseudovirus neutralization testing of matched serum was performed using a spike protein pseudotyped virus in 293/ACE2 cells, with neutralization assessed via reduction in luciferase activity (^32, 33^).

For binding ELISAs, matched serum was analyzed in duplicate on plates coated with SARS-CoV-2 NP or RBD (Zalgen Diagnostics, Aurora, CO) at 1:100 in diluent. Serum was incubated for 30 minutes at room temperature, washed four times, and incubated with anti-NHP IgG conjugate followed by incubation for 30 minutes at room temp.

Development by TMB for ten minutes was followed by stopping of the reaction and reading the plate at 450nm.

### Serum Cytokines

Invitrogen 37-Plex NHP ProcartaPlex kits were purchased and processed according to manufacturer’s instructions with a 1-hour sample incubation period and analysis on the Luminex xMAP. Heatmaps were generated using log_2_-transformed raw fluorescent intensity values input into the R package pheatmap (Raivo Kolde (2019). pheatmap: Pretty Heatmaps. R package version 1.0.12.). Hierarchical clustering was unsupervised.

### BAL Cytokines

Invitrogen 37-Plex NHP ProcartaPlex kits were purchased and processed according to manufacturer’s instructions with an overnight sample incubation period and fixation of the plate for one hour in 2% paraformaldehyde before resuspension in Reading Buffer and analysis using the Luminex xMAP. Heatmaps were generated using log_2_- transformed raw fluorescent intensity values input into the R package pheatmap (Raivo Kolde (2019). pheatmap: Pretty Heatmaps. R package version 1.0.12.). Hierarchical clustering was unsupervised.

### Pathology and Histopathology

Animals were humanely euthanized following terminal blood collection. The necropsy was performed routinely with collection of organs and tissues of interest in media, fresh frozen, and in fixative. The left and right lungs were imaged and then weighed individually. A postmortem bronchoalveolar lavage (BAL) was performed on the left lower lung lobe. Endoscopic bronchial brushes were used to sample the left and right mainstem bronchi. One section from each of the major left and right lung lobes (anterior, middle, and lower) sample fresh, and the remaining lung tissue was infused with fixative using a 50 mL syringe and saved in fixative. Fixed tissues were processed routinely, embedded in paraffin and cut in 5 µm sections. Sections were stained routinely with hematoxylin and eosin or left unstained for later analysis via fluorescent immunohistochemistry. Trichrome staining was performed as described previously, except with an additional 10 minutes of incubation with Weigert’s Iron Hematoxylin Working Solution^34^.

Histopathologic lesions identified in tissues were categorically scored by the same pathologist that performed the necropsies. Lesions were scored based on severity as the lesions being absent (-), minimal (+), mild (++), moderate (+++), or severe (++++). Cohorts were grouped together and non-parametric pairwise comparisons were performed for statistical analysis of histopathologic lesions.

Fluorescent immunohistochemistry was performed on 5 μm sections of Formalin-fixed, paraffin-embedded lung were incubated for 1 hour with the primary antibodies (SARS, Guinea Pig, (BEI, cat#NR-10361) diluted in NGS at a concentration of 1:1000). Secondary antibodies tagged with Alexa Fluor fluorochromes and diluted 1:1000 in NGS were incubated for 40 minutes. DAPI (4’,6-diamidino-2-phenylindole) was used to label the nuclei of each section. Slides were imaged with Zeiss Axio Scan Z.1 slide scanner. Other antibodies used for fluorescent immunohistochemistry are listed in Table S3.

### Quantification of pulmonary fibrosis

Trichrome stained slides were scanned on a digital slide scanner (S360, Hamamatsu Corporation, Bridgewater, NJ, USA). Annotation regions were drawn around the entire section of lung. The annotated regions were analyzed with a deep learning algorithm (HALO AI, Indica Labs, Albuquerque, NM, USA) trained by a pathologist (RVB) to recognize areas of fibrosis. The area of fibrosis was reported over the total area analyzed (% Area analyzed).

*Quantification of fluorescent immunohistochemistry for CD163+ and CD206+* Fluorescent immunohistochemistry was performed on sections of lung. Fluorescently labeled slides were scanned with a digital slide scanner (Axio Scan.Z1, Carl Zeiss Microscopy, White Plains, NY). Phenotypic markers were quantified using digital image analysis software (HighPlex Fl v4.04, HALO, Indica Labs). Cells were first identified by detecting nuclei (DAPI signal) and thresholds for detection of each phenotypic marker were set by a pathologist (RVB). The entire lung section was analyzed, and the number of each cellular phenotype (CD163+, CD206+, CD163+CD206+) was reported as cellsper mm^2^.

### Flow cytometry

After collection and processing of BAL as previously described, samples were centrifuged, cells resuspended in ammonium chloride potassium (ACK) lysis buffer (#A1049201, Fisher Scientific, USA), and incubated on ice. Media was added to stop lysis and cells were washed before counting and added to 5 mL snap-cap tubes. Cells were stimulated with 0.1% LPS (Enzo Life Sciences Cat# ALX-581-010-L001) and Brefeldin A (BD Bioscience Cat# 555029) overnight (16-18 hours). After stimulation, samples were washed with PBS pH7.2 (Fisher Cat#20012027) and incubated with Fc Block (Tonbo Biosciences Cat# 70-0161-U500) for 5 minutes on ice before viability staining (BD Biosciences Cat# 564406). Cells were washed with Running Buffer (Miltenyi Cat#130-091-221) before incubation with Surface Master Mix for 30 minutes on ice. After subsequent washing with Running Buffer, cells were resuspended in Fixation/Stabilization Buffer (BD Biosciences 554722) for one hour, then washed with Perm/Wash Buffer (BD Bioscience Cat# BDB554723) twice. Intracellular target antibodies were then added and incubated for 20 minutes at room temperature, then washed with Running Buffer. Samples were resuspended in FACS Fixation and Stabilization Buffer (BD Biosciences Cat# 50-620-051) and analyzed within 24 hours or resuspended in PBS and read between 48-72 hours after sample processing.

Compensation panels, pooled unstained sample, and unstimulated controls incubated with Brefeldin A only were run with every set of samples.

### Analysis of flow cytometry data

FlowJo version 10.7.1 (BD, Oregon, USA) was used to analyze flow cytometry data. Acquired samples were gated on viability, single cells, CD20/CD3 negativity, and HLA-DR positivity before cell typing. Total alveolar macrophages were gated based on their expression of HLA-DR, CD45, CD163, and CD206. Infiltrating macrophages were gated based on their expression of HLA-DR, CD45, and CD163 along with a lack of CD206 positivity. Monocyte-derived macrophages in the CD163+CD206+ population were distinguished from remaining alveolar macrophages by CCR2 and CD16 positivity.

Table S4 lists the antibodies used for staining, and a representative gating strategy (Figure S5).

### TGF-β ELISA

TGF-β1 was quantified in BAL supernatant fluid utilizing a Quantikine® ELISA TGF-β1 Immunoassay kit (R&D Systems #BD100B) according to manufacturer’s instructions and utilizing a Sample Activation Kit 1 (R&D Systems #DY010) and Human TGF-β1 Controls (R&D Systems #QC01-1). Each animal’s necropsy BAL supernatant was compared to baseline samples, and log^2^ fold change was calculated.

### Hematology and Clinical Chemistries

Analysis of blood chemistries was performed using a Sysmex XT-2000i analyzer for EDTA collected plasma, or an Olympus AU400 chemistry analyzer for serum.

## Results

### Viral Dynamics

Eight rhesus macaques (RMs) and eight African green monkeys (AGMs) were inoculated with SARS-CoV-2, with a subset (n=4) of each species experimentally infected either by multiroute (IT/IN) or small particle aerosol. The individuals received 2.0 x 10^6^ TCID_50_ via IT/IN and approximately 1.5 x 10^4^ TCID_50_ via aerosol, ranging from 1.9 x 10^3^ to 7.5 x 10^4^, depending on each animal’s respiratory patterns (Table S1). Mucosal sampling via pharyngeal, nasal, and rectal swabs, blood, BAL and radiography were performed at indicated times (Figure 1).

Quantitative RT-PCR was used to measure viral load of both genomic and subgenomic vRNA throughout the course of disease (Table S2). Viral loads were generally at their peak at one day post challenge, with viral RNA trailing off for most species/routes by the end of week 2. All cohorts, except the RM aerosol cohort, had detectable genomic RNA at necropsy in the nasal swabs, though nothing was still detectable in the pharyngeal swab. In the BAL supernatant, genomic RNA persisted longer in the aerosol groups as compared to their species-matched IT/IN exposure cohort, with the AGM aerosol cohort still having detectable genomic vRNA at necropsy, and detectable subgenomic vRNA two weeks post challenge. Persistent, delayed shedding of viral RNA was observed in rectal swabs of the aerosol cohorts across species, with subgenomic vRNA present at necropsy in the AGM aerosol animals (Figure 2A). Overall viral loads, represented as area under the curve of vRNA over the course of the study, overall higher genomic vRNA in the multiroute compared to the aerosol cohort in the RMs, with the same relationship being present in rectal swabs for subgenomic vRNA (Figure S1).

**Figure 2:**
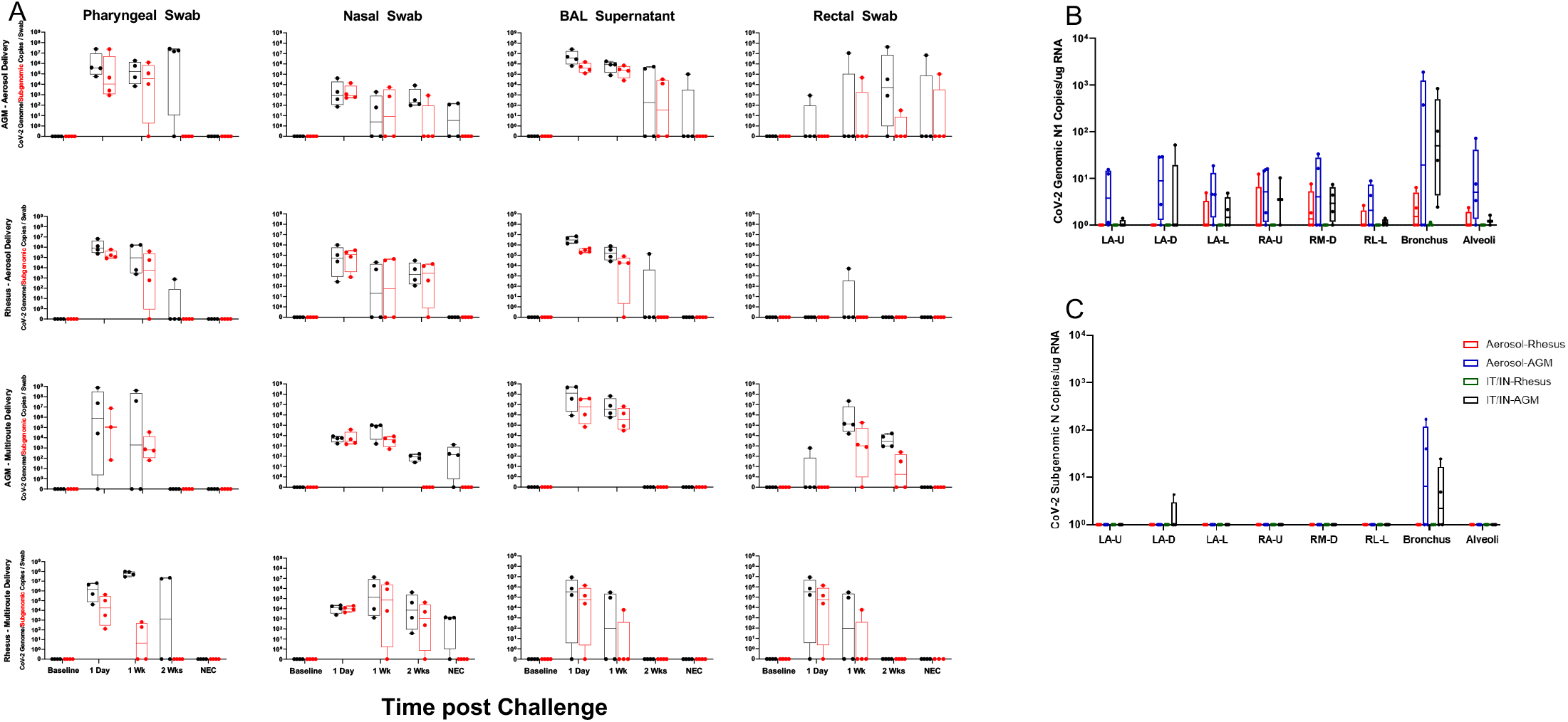
Viral Loads Assessed via RT-qPCR post SARS-CoV-2 Challenge. A)Viral loads in swabs and BAL supernatant were assessed via RT-qPCR post challenge for genomic (black) and subgenomic N (red) RNA. After necropsy, respiratory tissues were analyzed for the presence of genomic (B) and subgenomic (C) content.

Genomic RNA was detected in multiple lung regions, with the notable exception of the RM multiroute cohort, where no persistent RNA was found at necropsy (Figure 2B). All AGM animals displayed subgenomic vRNA in lung tissue at necropsy, with detectable amounts of viral RNA in the bronchus and one individual maintaining detectable amounts in the LA-D region (Figure 2C).

Live virus was measured in each sample via the median tissue culture infectious dose (TCID_50_) assay. Viral loads followed a similar pattern of a high peak at one day post challenge, in all animals regardless of exposure modality or species. The aerosol cohort exhibited longer term viral loads, with one AGM still possessing detectable virus at week three post challenge in pharyngeal swab samples. Virus was detected in nasal swabs and BAL supernatant of the AGMs regardless of exposure, with much less detected in the RMs (Figure S2).

### Antibody Responses

We used a combination of enzyme-linked immunosorbent assay (ELISA) and pseudovirus neutralization to characterize the antibody responses of the cohorts during SARS-CoV-2 infection. In all cases, with pseudovirus neutralization (Figure 3A), and binding of RBD and NP (Figures 3B and C, respectively), RMs showed less antibody development than AGMs, regardless of exposure modality, though their anti-RBD development resembled the AGM aerosol cohort (Figure 3B). The RM aerosol cohort displayed anti-NP titers equivalent to the AGM aerosol animals at week three, with a drop in titers at necropsy (Figure 3C). The AGM mutiroute cohort showed the highest magnitude of antibody development, with a peak at week 3, then a slight decrease from peak by necropsy. Notably, the AGM aerosol cohort displayed an overall delayed antibody response, which peaked at necropsy, indicating the potential to continue to increase at later time points. This peak in titers was observed in AGMs in both exposure cohorts (Figure 3D). In all cases, a higher titer was seen at necropsy in the AGM cohort regardless of exposure modality than the RM cohort, with no difference observed based on exposure modality (Figure 3E-H).

**Figure 3:**
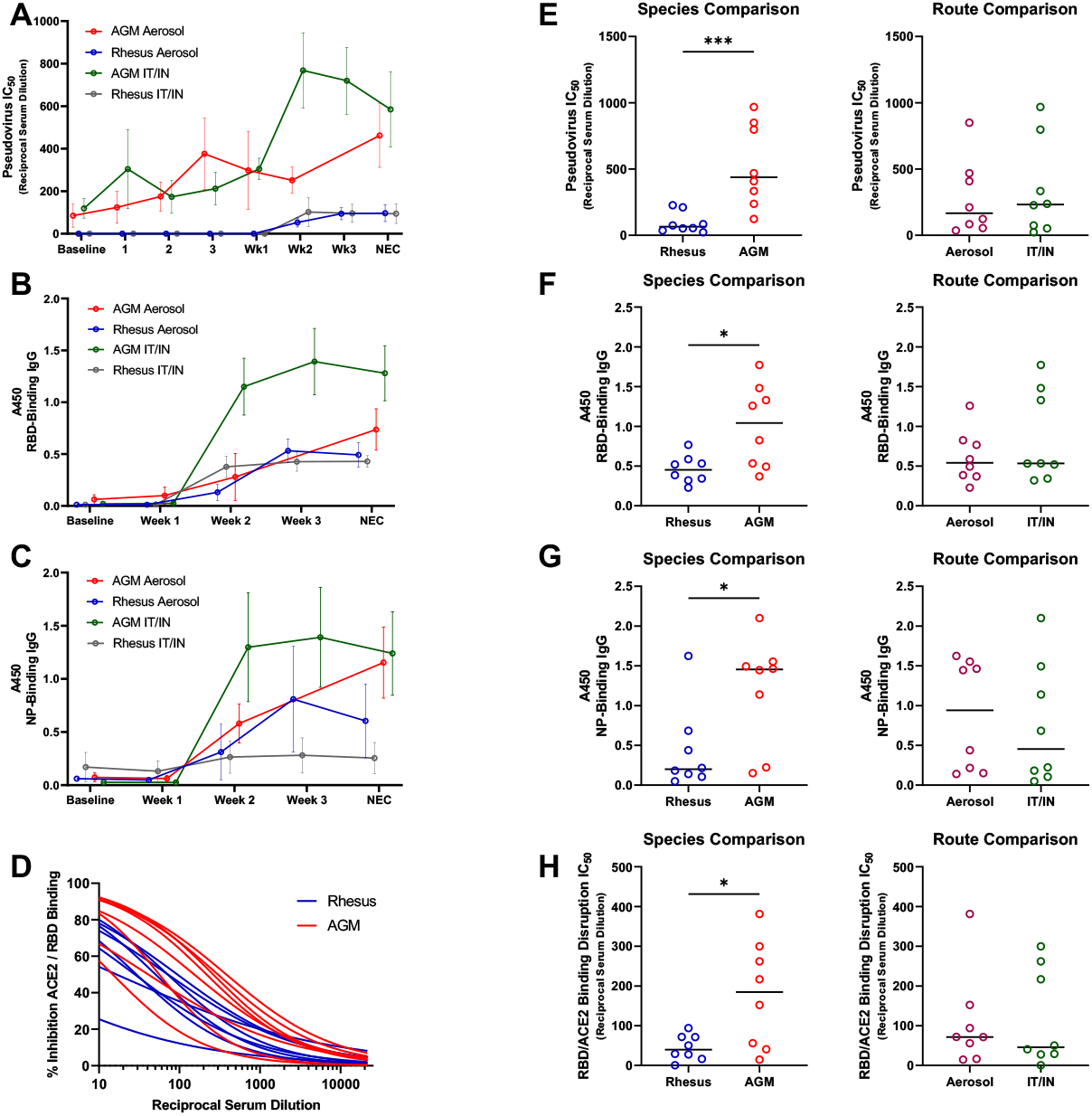
Antibody Responses to SARS-CoV-2 Challenge. Antibody responses were followed post challenge by A)pseudovirus neutralization, B and C) ELISA for binding to RBD and NP, respectively, and D) surrogate virus neutralization test at necropsy. Route and species variability in antibody levels at time of necropsy were compared for each assay type (E-H). Comparisons were made using the Mann- Whitney or Welch’s t-test, depending on normality of data. Asterisks represent significant comparisons (*, p<0.05; ***, p<0.001).

### BAL and Serum Cytokines

Measurements of cytokines in serum and BAL supernatant samples were performed throughout the study. Cytokines in the BAL supernatant was more differentially expressed than those in the serum. BAL cytokines were expressed in higher amounts in the mutiroute than the aerosol cohort, with the aerosol animals revealing a delayed increase in many cytokines, including IL-2, IL-4, IL-6, IL-7, IL-10, TNF-α, and IFN-α. The RM multiroute cohort exhibited a larger increase in some cytokines than the AGM cohort, including GM-CSF, IL-10, IL-4, and IP-10, and an increase above the RM aerosol animals in IL-12p70, IL-17A, TNF-α and IL-7. The aerosol cohort exhibited lower levels of MCP-1, with the aerosol-exposed RMs expressing IP-10 more highly than the AGM aerosol animals. In some cases, aerosol continued to trend at necropsy with increases of IL-6, IFN-α, IL-1β throughout sampling. AGMs displayed higher levels of I-TAC and lower levels of MCP-1 than RMs, though trends were more associated with exposure modality than species in the BAL (Figure 4).

**Figure 4:**
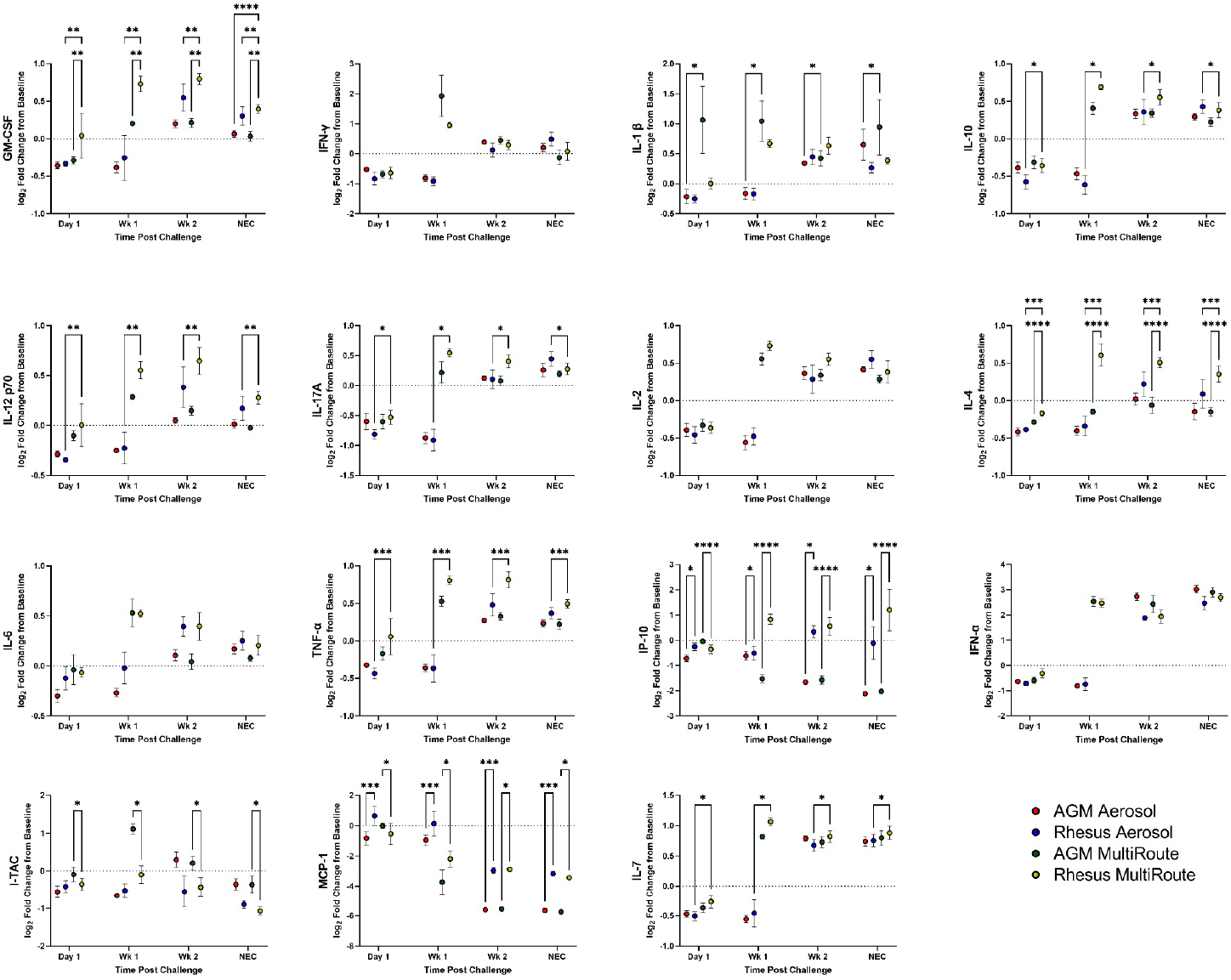
BAL Supernatant Cytokines post SARS-CoV-2 Challenge. Cytokines in BAL supernatant were analyzed at indicated time points post challenge. Comparisons were made with two-way ANOVA using Tukey’s multiple comparisons test. Asterisks represent significant comparisons (*, p<0.05; **, p<0.01;***, p<0.001;****, p<0.0001).

Serum cytokines were measured throughout the study as well. Here, the RM multiroute animals displayed lower levels of GM-CSF, IL-5, IL-6, and MCP-1. The temporal differences in exposure modality were not present in the serum samples. The AGM mutiroute displayed higher levels of MIG, MIP-β1 and VEGF-A (Figure S3).

### Clinical Scoring

Clinical observations were performed throughout the study and resulted in categorical scores as a corollary to clinical disease development in the NHPs. The number of animals showing signs of SARS-CoV-2 related disease increased rapidly early during the study in the mutiroute cohort in either species, continued to a peak at week three post challenge, and then declined by day 28 termination of the experiment (necropsy).

This is in contrast to the aerosol-exposed animals, which did not display signs of disease until one week post challenge, and thereafter continued to increase until day 28 at necropsy (Figure 5A, B). The same pattern was observed in overall group severity scores (Figure 5C). Overlaying the clinical score curves with PCR data showed persistent subgenomic vRNA in the BAL supernatant (Figure 5A) and rectal swabs (Figure 5B) of the AGM aerosol cohort, suggesting a slower onset, more persistent disease process. Both species exhibit persistent signs of disease, with RMs peaking than AGMs at necropsy over that of the AGM animals (Figure 5D) when data is segregated by species among all of the aerosol-exposed animals. Overall, the mutiroute cohort exhibited a higher cumulative score than the aerosol cohort, though the latter cohort’s clinical signs continued to increase until the termination of the study (Figure 5E). The scoring system was based on respiratory signs and changes in activity (Figure 5F).

**Figure 5:**
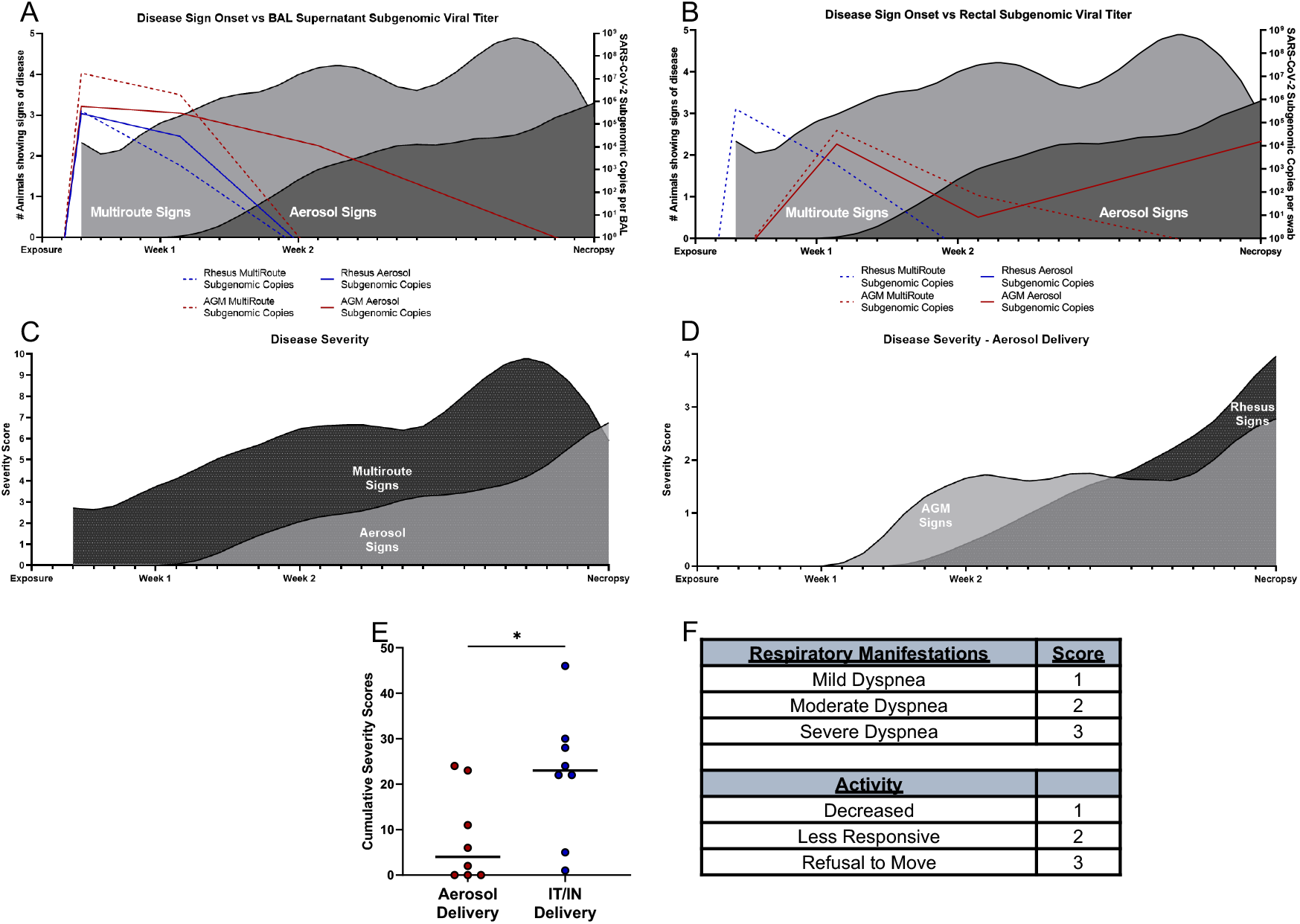
Clinical Disease Severity Scores. Signs of disease present in each cohort over time. Number of animals showing signs of disease along with subgenomic viral loads in A) BAL supernatant and B) rectal swabs. Severity scores per group in C) both delivery cohorts and D) aerosol cohort split into species. E) Cumulative severity scores per delivery cohort. F) Simplified scoring system used for cohorts. Curves in figures were smoothed. Comparisons made using Welch’s t-test. Asterisks represent significant comparisons (*, p<0.05).

### Pulmonary Fibrosis

Ten animals of the sixteen exposed to SARS-CoV-2 infection by either aerosol or mutiroute presented with pulmonary scarring of varying but generally mild degree, including animals with no signs of disease observed. To define animals with and without pulmonary scarring for further analysis, three criteria were applied as compared to a normal control (Figure 6A): fibrin deposition observed by a licensed pathologist using routine H&E histology (Figure 6B); trichrome staining to identify areas of inflammation and collagen deposition in at least one section of lung (Figure 6C); and analysis with HALO, per indicated methods, revealed at least one section of lung which contained more collagen than one standard deviation above the average identified in the same section in naive animals (Figure 6D). Animals classified with pulmonary scarring were generally identified based on sections collected from the deep left anterior lobe section.

**Figure 6:**
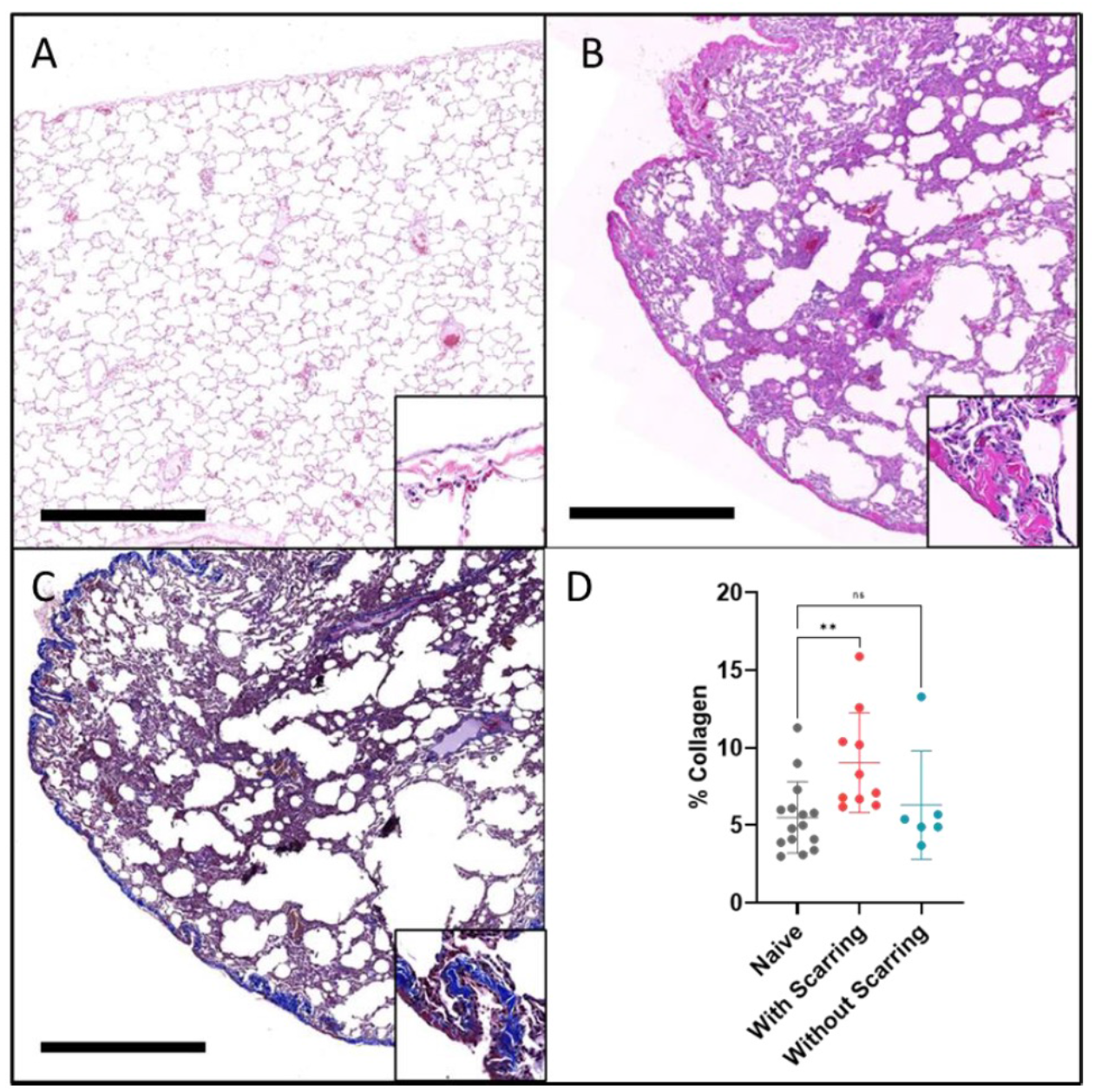
Mild pulmonary scarring in NHPs exposed to SARS-CoV-2. Representative figure of sequentially cut slides of lung tissue from a SARS-CoV-2- infected NHP. A) H&E of normal lung tissue. Inset: normal pleura characterized by an outer mesothelial lining and a thick layer of collagen. B,C: KN90, LA-D. There is mild, diffuse thickening of the collagen layer of the pleura on B) H&E and C) trichrome, blue) stains. Insets show higher magnification of the pleural fibrosis. Bar = 1mm. D) Percent collagen identified in trichrome stained lung by HALO analysis. **p<0.002 , one-way ANOVA with Dunnett’s multiple testing correction.

### Lung Macrophage Dynamics

Flow cytometric analysis of CD163+CD206+ alveolar macrophages in BAL fluid revealed an average decrease in this population at day one post-exposure in animals with and without pulmonary scarring, with recovery to naïve comparator levels in animals with scarring over the course of the study (Figure 7A). Infiltrating macrophages within the lumen of the alveoli, classified as CD163+CD206-, were higher at day one than in naïve comparators, and resolved by week one in animals with scarring.

**Figure 7:**
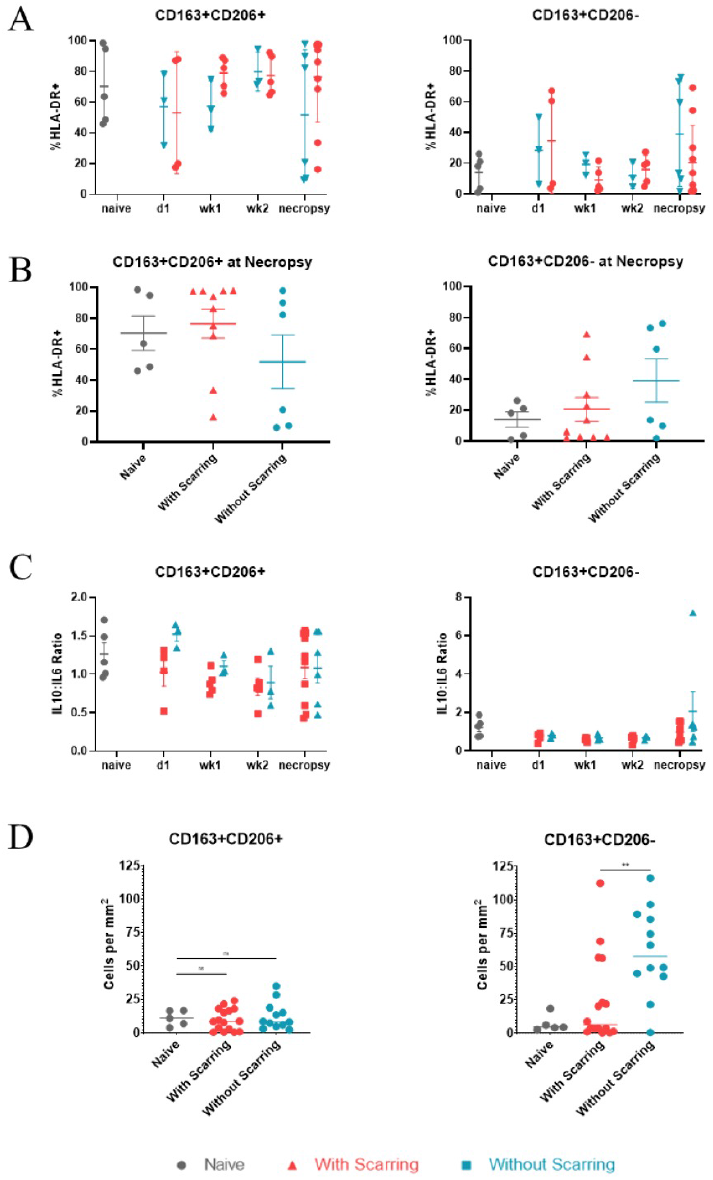
Myeloid cell kinetics in BAL. A) Quantification of alveolar macrophage (CD163+CD206+) and infiltrating macrophage (CD163+CD206-) populations throughout the study. B) Quantification of alveolar and infiltrating macrophages at necropsy. C) IL- 10:IL-6 ratio calculated from mean fluorescence intensity (MFI); Right, Pearson correlation test p=0.001. D) HALO analysis of macrophage phenotypes reported as cells per mm^2^.

Meanwhile, infiltrating macrophages remained high in animals without scarring (Figure 7A). Animals with scarring had more alveolar macrophages and fewer infiltrating macrophages than animals without scarring at necropsy (Figure 7B). IL-10:IL-6 ratios were calculated via quantification of median fluorescence intensity (MFI) and revealed greater IL-10:IL-6 ratios in both alveolar and infiltrating macrophages in animals without pulmonary scarring (Figure 7C). Additionally, HALO analysis was completed with fluorescent markers for both CD206 and CD163 to validate flow cytometry findings and confirmed increased macrophage presence in pulmonary tissue (Figure 7D). Combined quantification and functionality analysis indicated increased anti-inflammatory macrophages in the pulmonary compartment of animals without pulmonary scarring.

Further analysis of alveolar macrophages (CD163+CD206+) within the lung included distinction between CD163+CD206+CD16+CCR2+ infiltrating monocyte-derived alveolar macrophages and CD163+CD206+CD16- resident alveolar macrophages. The proportion of alveolar macrophages decreased in both groups at day one and remained low through the end of the study, while monocyte-derived macrophages dominated the CD206+CD163+ subset (Figure 8A). Both alveolar macrophages and monocyte-derived macrophages were higher in animals without pulmonary scarring and produced higher IL-10:IL-6 ratios throughout the study (Figure 8B), suggesting a more robust, persistent, and anti-inflammatory pulmonary compartment in animals without scarring.

**Figure 8:**
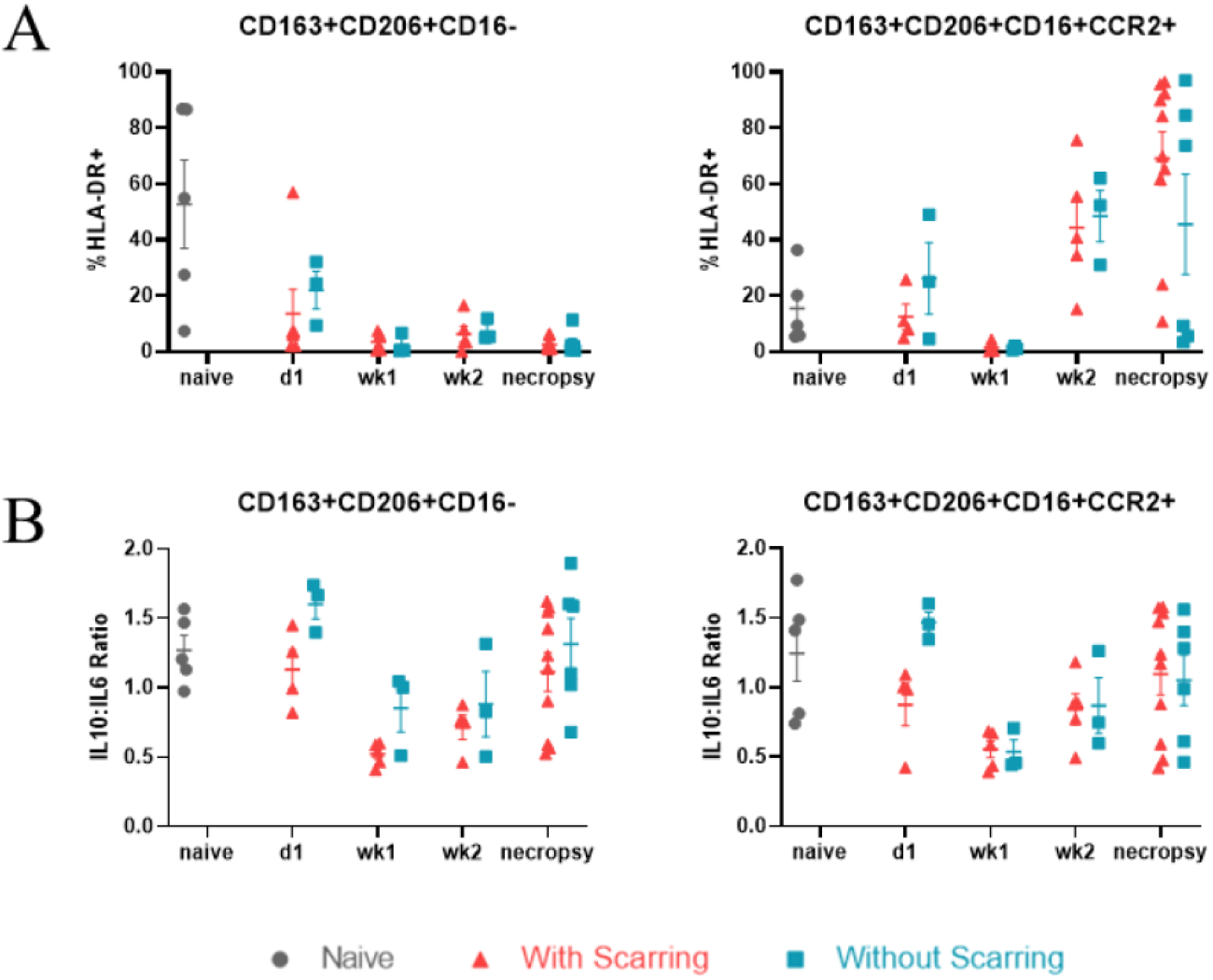
Infiltration and persistence of infiltrating macrophage. A) Left: Resident alveolar macrophages (CD163+CD206+CD16-) as a percent of alveolar macrophages; p=0.003, ordinary one-way ANOVA. Right: Monocyte-derived alveolar macrophages (CD163+CD206+CD16+CCR2+) as a percent of alveolar macrophages. B) Ratio of IL- 10:IL-6 in Left: resident alveolar macrophages, Right: monocyte-derived alveolar macrophages; calculated from MFI.

### Long-term myofibroblast persistence and lack of regenerative activity in areas of pulmonary fibrosis

Pulmonary scarring is the result of a combination of both persistent fibroblast activation and reduced regenerative potential, causing excessive deposition of collagen within the lung that is not readily resolved. To determine if activated myofibroblasts were present within this collagenous tissue, lung sections of animals both with and without pulmonary scarring were stained with fluorescent markers for αSMA and cytokeratin V. Myofibroblasts were present in areas of collagen deposition (Figure 9A). Myofibroblasts are activated by TGF-β, which is produced by macrophages and allows for their continued production and deposition of collagen. Additionally, TGF-β signaling has also been implicated in fibrogenesis^35, 36^. To determine if activated myofibroblasts were within collagenous lung sections, we measured for the presence TGF-β in BAL fluid.

**Figure 9:**
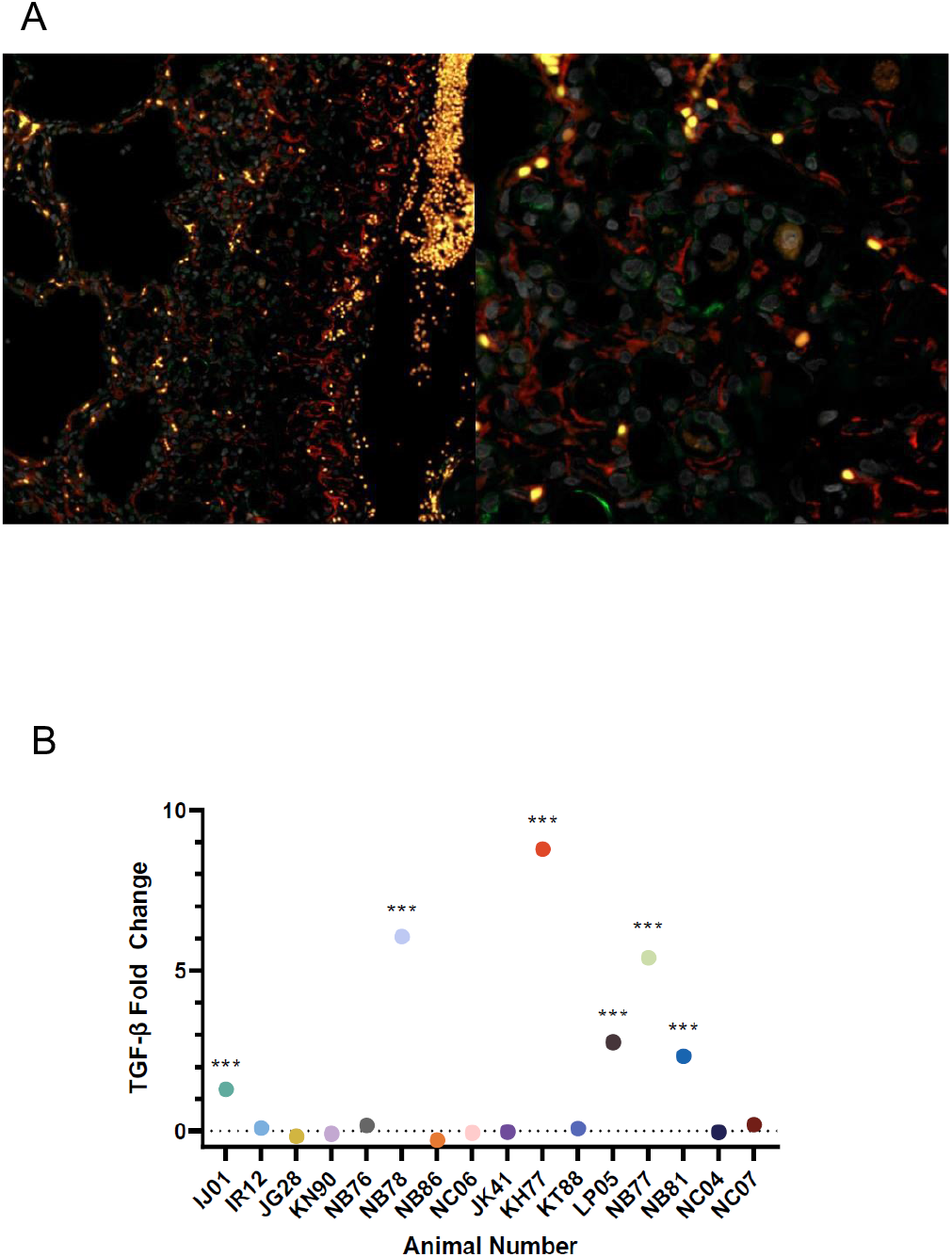
Activated myofibroblasts are present within scarred lung tissue and persist long term. A) KN90, lung alveoli. Myofibroblasts characterized by double-positive staining of αSMA (red) and cytokeratin V (green). Bar = 100μm. B) Continued myofibroblast activation, as determined by TGF beta ELISA, at 28 days post exposure. ***p<0.001, multiple unpaired t-test with Holm-Šídák correction.

Detectable levels of TGF-β were in some, but not all, of animals with lung scarring, and in none of the animals without lung scarring already present (Figure 9B). The presence of TGF-β was not discernable between species nor modality of SARS-CoV-2 infection. Because TGF-β is closely involved in wound healing and associated immunoregulation at sites of injury, there was interest in understanding whether regenerative processes were taking place in animals recovering from acute viral infection. Fluorescent antibodies p63 and cytokeratin V were used to identify basal lung progenitor cells to measure whether regenerative potential within the lung was underway. We identified a lack of robust regenerative response to SARS-CoV-2 infection (Figure 10A-D), which is suggestive of a muted regenerative response and potentially prolonged recovery from viral infection in the lung of experimentally infected macaques.

**Figure 10:**
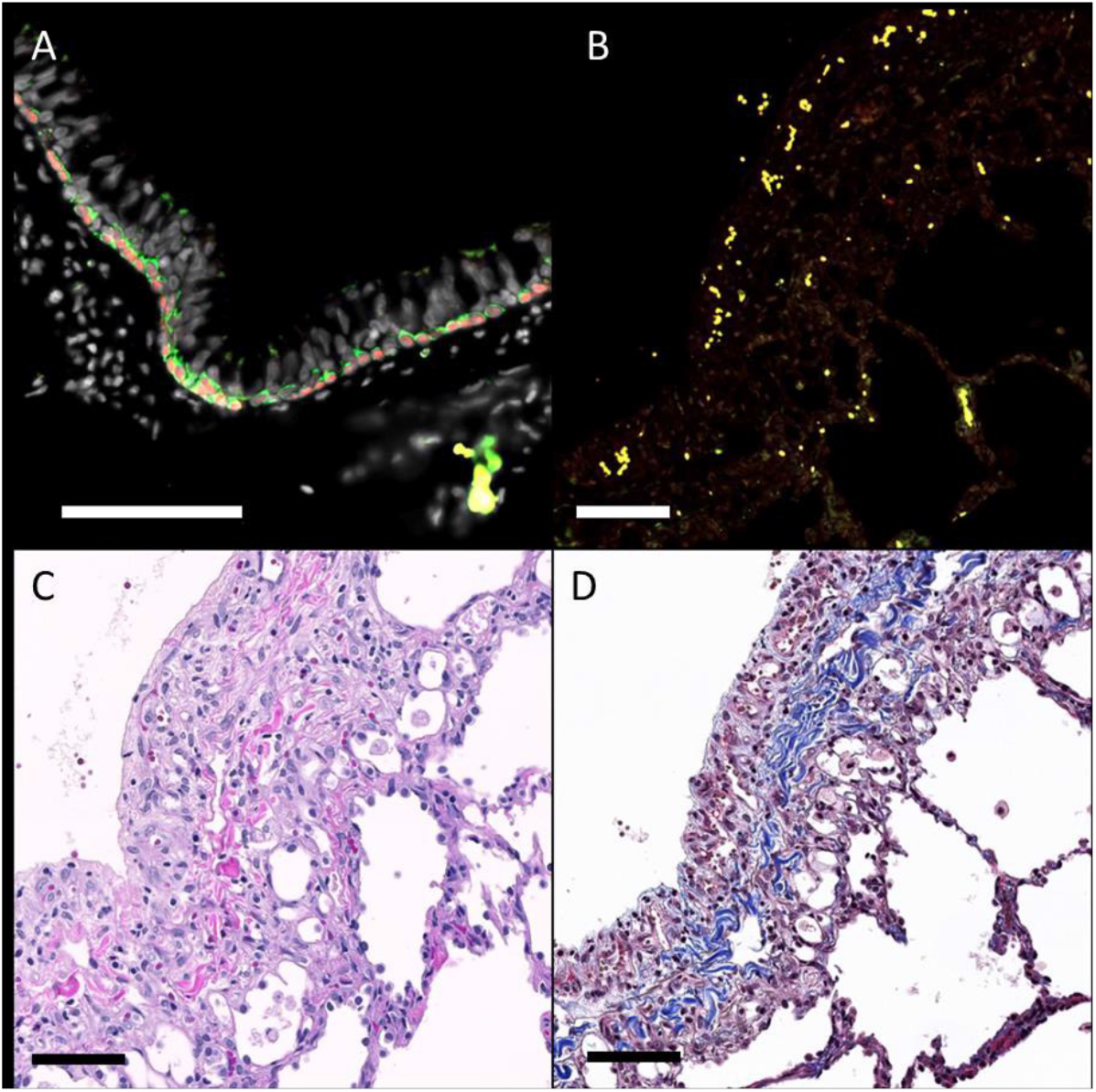
Absence of regenerative activity at 28 days post-exposure. Immunofluorescence for the detection of bronchial epithelial progenitor cells. A) NJ48, lung. The epithelial lining of large airways (bronchi) has a basal layer containing progenitor cells characterized by cytoplasmic expression of cytokeratin V (green) and nuclear expression of p63 (red). B-D) KN90, lung alveoli. B) Progenitor cells are not observed in regions of pleural fibrosis, even in regions where moderate fibrosis and inflammation are observed with H&E (C) and trichrome (D). Bar = 100 μm.

### Hematology and Clinical Chemistries

Complete blood counts and clinical chemistries were performed throughout the study at times of blood collection. The AGM mutiroute cohort displayed lower levels of neutrophils than the AGM aerosol animals during the study (Figure S6C) and higher levels of monocytes than the AGM aerosol and RM multiroute cohorts (Figure S6F).

Blood urea nitrogen was elevated in the AGM aerosol cohort compared to the RM aerosol cohort (Figure S7D), though overall increases in BUN were only slightly higher than expected levels for either NHP species.

## Discussion

SARS-CoV-2 infection continue to result in morbidity and mortality in large segments of the population, with some long-term sequelae persisting well beyond viral clearance in those that carry risk factors associated with severe clinical outcome^1, 37–41^. Variants of concern (VOC) are arising that threaten to continue the pandemic into the future to an uncertain degree, necessitating efforts that transcend vaccination strategies alone, and promote alternative prophylactic and therapeutic development both in acute and longer-term time frames post-infection^42–46^. Multifocal medical countermeasure development while this virus evolves to the human host necessitates a suitable animal model of infection that recapitulates the immunologic and clinically relevant aspects of COVID-19. Accordingly, we evaluated both the multiroute and aerosol exposure modality of SARS-CoV-2 experimental infection in the Rhesus macaque and African green monkey. In this study, we attempted to dose animals using two distinct exposure routes: intratracheal and intranasal concomitantly, and by small particle aerosol. The dosing in the ‘mutiroute’ groups were fairly easy to obtain as doses were tittered based upon volume applied in the administration in both the intratracheal and intranasal administration. Reaching the equivalent dose (2E+06 TCID_50_) by the aerosol route was hampered by relatively low-titer virus (3E+06 TCID_50_) and the logistics of producing aerosols for the purposes of individual exposure. There is a natural dilution effect and corresponding efficiency to aerosolization of SARS-CoV-2 virus that effects the resulting dosing of each animal. The net effect was a nearly 2-log disparity between the multiroute and aerosol groups. This disparity should be considered when any direct comparisons are made controlling only for route of exposure.

Animals in the mutiroute exposure group displayed earlier and respiratory signs of disease of increased severity, as well as a higher cumulative categorical clinical score than the aerosol cohort. The aerosol-exposed animals of either species, in contrast, began showing signs of disease a full week later and continued to trend upward until the termination (28d PI) of the study. Though signs of disease were delayed in the aerosol cohort, viral titers were similar beginning day one through week one post-exposure in all in all groups, with persistent infection detected in aerosol exposure cohorts through end of the study. The similarity of the pattern of early viral kinetics between exposure groups despite differences in clinical signs of disease suggests that viral replication may not be the direct cause of disease onset, but rather the immune response to infection that may play a larger role. Additionally, animals who did not display signs of disease throughout the study nonetheless exhibited detectable measures of viral replication, including PCR detection of viral genomic and subgenomic RNA, indicating that these animals may still be able to transmit virus. Higher viral titers in the lower respiratory tract were observed in AGMs, whereas the nasal cavity positivity was greater in RMs, representing potential differences in natural aerosol transmission between species.

Systemic inflammatory cytokine/chemokine response including TNFα, IL-6, MCP-1, IP-10, and MIP-1α has been associated with more severe disease in human studies of COVID-19^3, 47^. We observed increases in some of these mediators early in infection in serum including MCP-1 and MIP-1α in AGMs of both exposure routes, indicating mobilization and recruitment of monocytes, dendritic cells, and NK cells in these animals. A single animal (NB81) in the AGM mutiroute group exhibited acute increases in IFN-γ, MIG, MIP-1β, and VEGF-A, and drove group trends in blood monocyte number. These outcomes indicate a contribution of NK cells to the activation of macrophages in this animal, as well as robust extravasation of monocytes.

Localized soluble mediator responses varied among groups and species. More pronounced changes were observed in the BAL than in the serum, indicating a more localized pulmonary than systemic response to infection in this model. The mutiroute group in both species showed more pronounced changes than the aerosol cohort, possibly due to dose differences. Similar increases in IL-2, IL-4, IL-6, IL-7, IL-10, TNFα, and IFNγ were observed at week one in mutiroute exposure groups, though the aerosol group saw similar increases a week later. This shift in cytokine response correlates with the shift observed with signs of disease, indicating a delayed immune response to infection despite temporally similar viral titers.

We found more alveolar macrophages and fewer infiltrating macrophages at nearly all time points by flow cytometry of BAL and at necropsy by HALO analysis of fluorescent antibody-stained lung tissue, in animals with pulmonary scarring. Infiltrating macrophages were increased in animals without scarring at necropsy, though their average IL-10:IL-6 ratio was higher than those in animals with scarring, indicating an anti-inflammatory phenotype. This finding emphasizes the need for further investigation into the role of inflammatory/anti-inflammatory phenotype of infiltrating macrophages in pulmonary scarring post-SARS-CoV-2 infection. Indeed, individual animals without scarring with fewer infiltrating macrophages had the highest IL-10:IL-6 ratios, alluding to a more potent anti-inflammatory response of individual cells in these animals. Individual animals with scarring and fewer infiltrating macrophages also had the lowest IL-10:IL-6 ratios, indicating a more potent inflammatory response. The numbers of both resident alveolar macrophages (CD163+CD206+CD16-) and monocyte-derived macrophages (CD163+CD206+CD16+CCR2+) are also higher in animals without scarring early in infection at day one, with a consistently higher average IL-10:IL-6 ratio in these cells.

Correspondingly, at day one PI, a drastic decrease in IL-10:IL-6 ratio in all alveolar macrophage (CD163+CD206+) populations in animals with scarring suggests a marked and early response, as seen in human cases with worsening disease^48^, sets the stage for separation between scarring outcomes. This early indicator of potential long-term consequences of disease merits further investigation and suggests that inflammatory/anti-inflammatory phenotype and potency of response of macrophages in the lung may be an important indicator to predict scarring outcome. This additionally suggests differences in proinflammatory monocyte-derived macrophage populations at baseline may dictate response, as well as the utility of anti-inflammatory steroids such as dexamethasone to prevent pulmonary scarring in SARS-CoV-2 infection by reducing the inflammatory response. SARS-CoV-2 results in death of monocyte-derived macrophages while still correlating with IL-6 secretion, which could result in this IL-10 dominated infiltrating macrophage response as observed in this study^49^. Focused studies thouroughly interrogating host response in this context should be performed based upon these findings.

Activated myofibroblasts positive for αSMA and cytokeratin V in areas of collagen deposition within scarred lung were identified using fluorescent immunohistochemistry.

Myofibroblasts co-localized with macrophages were identified within the pulmonary region of the lung, further suggesting their role in the activation and persistence within scarred tissue. The presence of activated myofibroblasts within scarred areas of tissue also suggests further development of this cellular subset in affected areas within the lung. Additionally, when TGF-β ELISA was performed on BAL fluid, a cytokine which induces activation of myofibroblasts and is associated with wound healing and found significant levels of TGF-β in some, but not all, animals with lung scarring at 28 days post-exposure. Further studies should explore the secretion and resolution of TGF-β production and its relation to pulmonary scarring after SARS-CoV-2 infection. Immunohistochemistry identifying basal progenitor cells positive for p63 and cytokeratin V showed minimal difference in the frequency of progenitor cells between animals naïve to SARS-CoV-2 and infected animals with or without pulmonary scarring after infection. This lack of signs of robust regenerative activity after infection suggests that infection with SARS-CoV-2 negatively affects the potential for prolonged recovery and resolution of pulmonary scarring in NHP. This finding is consistent with clinical human findings of elevated TGF- β in severe COVID^50^, as well as myofibroblast activity within infected lungs ^51^. This makes the NHP model of SARS-CoV-2 infection a powerfully predictive tool for this unique pathologic sequalae, as well as for testing future therapeutics that targets TGF-β^52, 53^.

Delayed local pulmonary cytokine/chemokine responses, prolonged viral titers through necropsy, and residual pulmonary inflammation at necropsy may indicate that aerosol inoculation produces prolonged consequences of infection. Further, broad dose differences (nearly 2-log) between mutiroute and aerosol groups produce similar outcomes, which suggests aerosol exposure is a more potent exposure modality for SARS-CoV-2 infection. The aerosol AGM group also showed consistent, modest signs of disease trending upward at necropsy, persistent viral titers in the BAL, and a trend of increasing inflammatory cytokines at necropsy, indicating aerosol exposure of AGMs as an appropriate NHP model of post-COVID syndrome.

Both aerosol RM and mutiroute AGM groups demonstrate increased respiratory rate, comparable neutrophilia, and increased signs of disease in their respective exposure group. However, the AGM mutiroute group exhibited the greatest lymphocyte and monocyte increases, earliest and greatest signs of disease, highest pharyngeal peak viral titer, most inflammatory cytokines in serum, and most consistently high BAL inflammatory cytokines (IL-6, IFNα, TNFα, IL-1β). Taken together, these responses suggest mutiroute exposure of AGMs most accurately recapitulates human disease with predictive poorer clinical outcome. Future studies should consider use of either species, and choice of exposure modality when exploring currently-identifed or future VOC as dictated by the tempo and sustainability of the COVID-19 pandemic.

---

**Figure S1:**
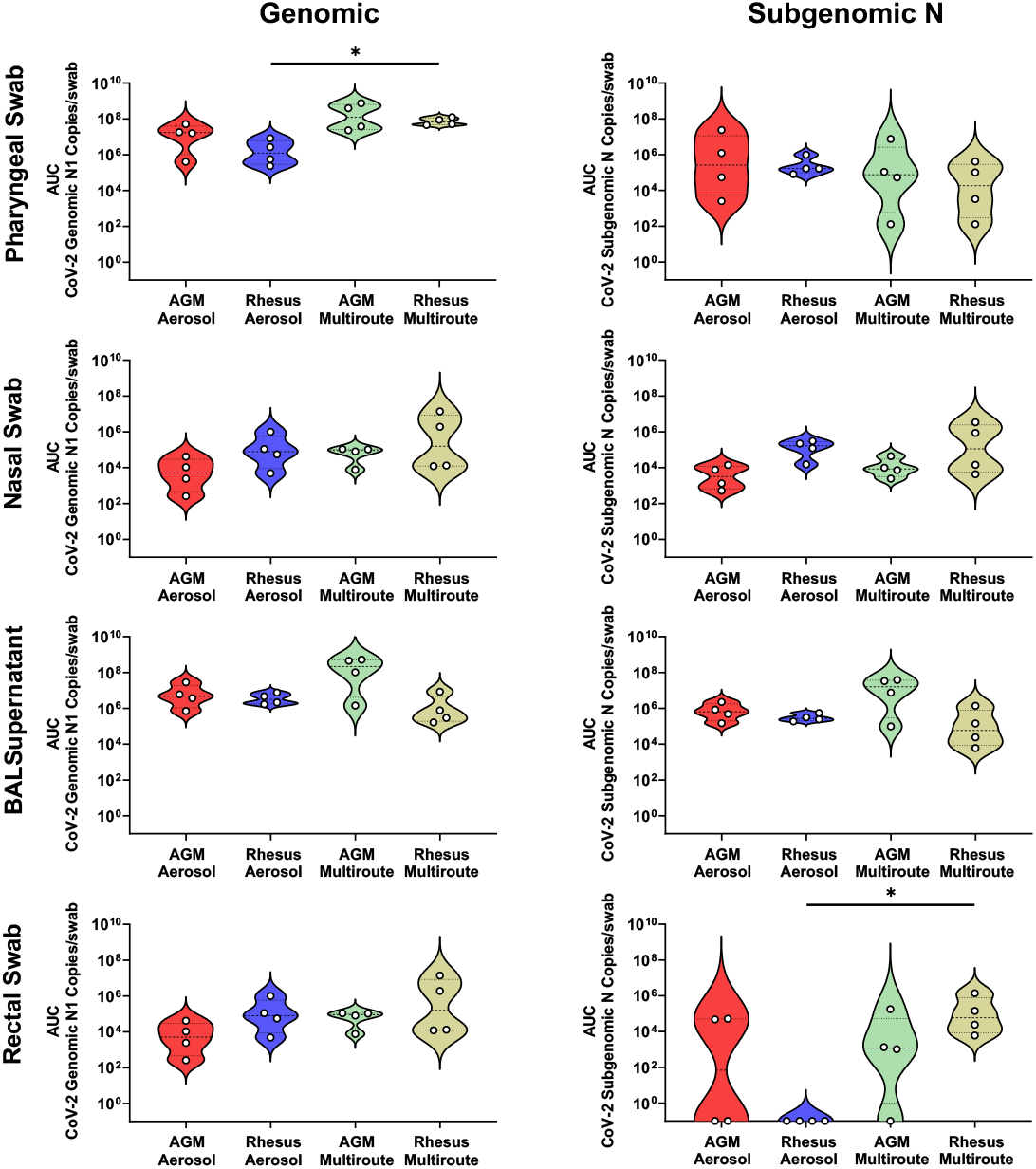
Viral Loads Assessed via RT-qPCR post SARS-CoV-2 Challenge. Viral loads, assessed by RT-qPCR for genomic and subgenomic RNA, represented as area under the curve for the post challenge period. Comparisons between groups were made via Kruskal-Wallis with Dunn’s multiple comparisons test. Asterisks represent significant comparisons (*, p<0.05).

**Figure S2:**
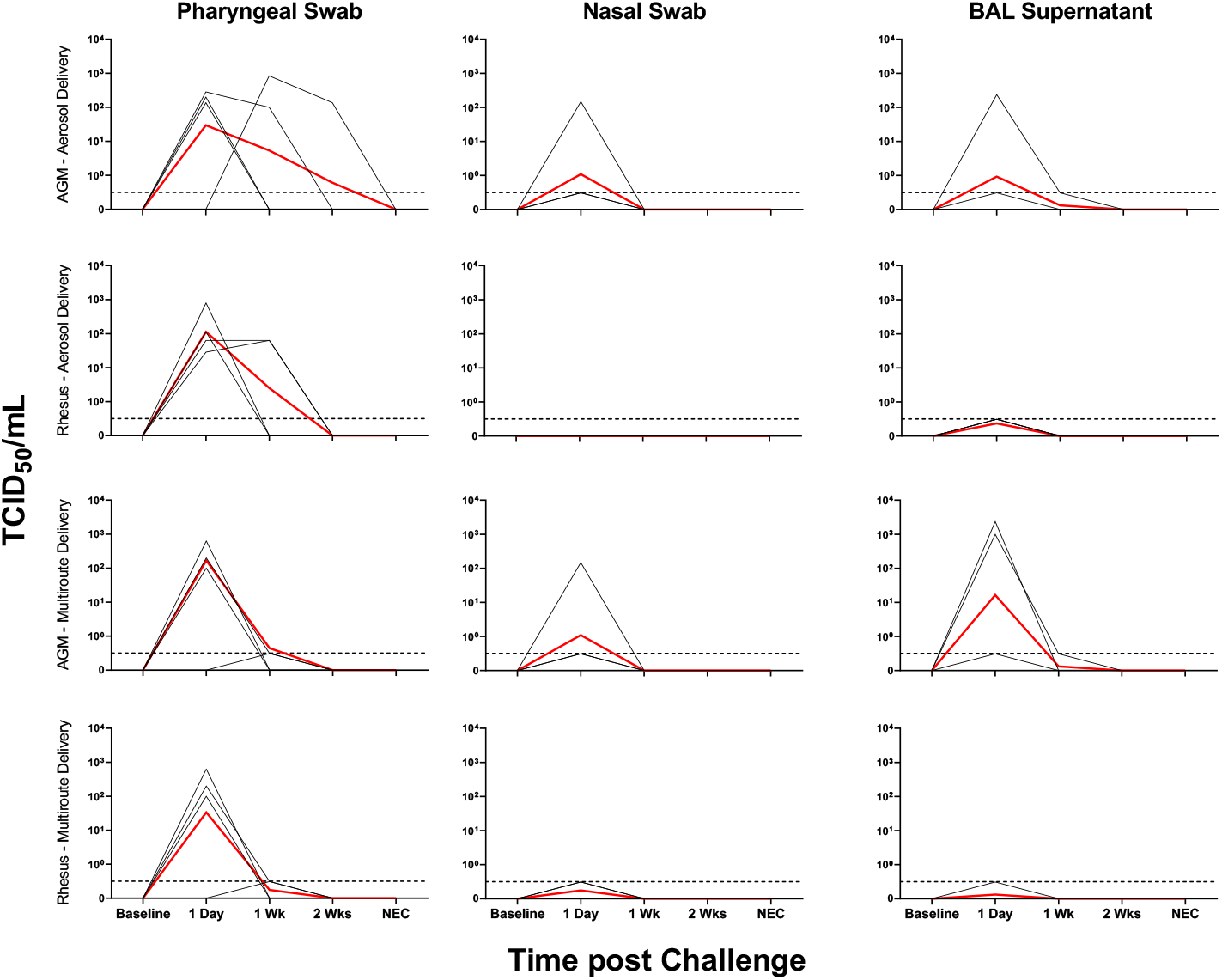
Viral Loads Assessed via TCID_50_ post SARS-CoV-2 Challenge. Viral loads, assessed by TCID_50_, represented as area under the curve for the post challenge period. Black lines indicate viral loads per individual, with red lines indicating group geometric means. Dotted lines indicate a positive sample below the limit of quantification.

**Figure S3:**
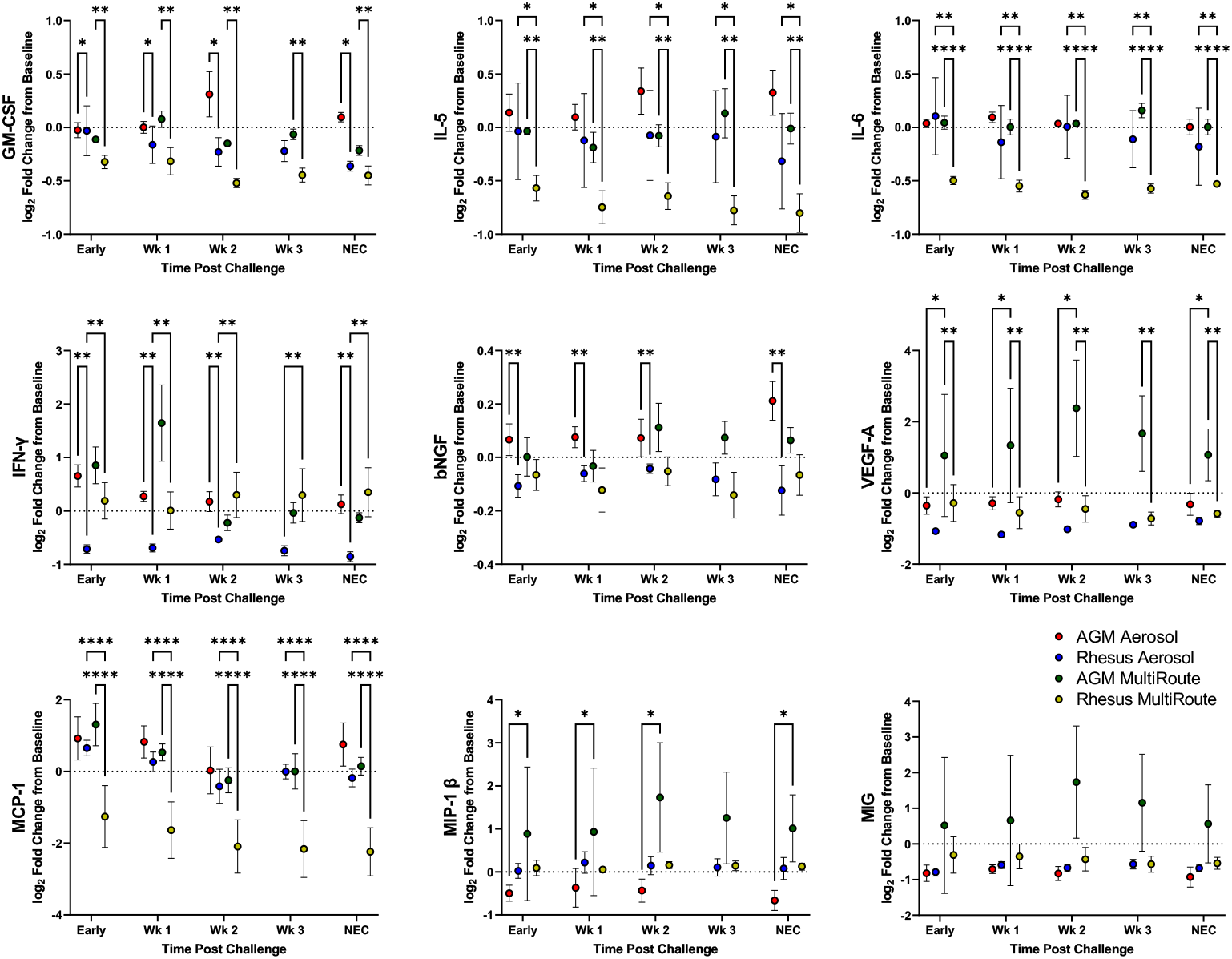
Serum Cytokines post SARS-CoV-2 Challenge. Cytokines circulating in serum were analyzed at indicated time points post challenge, with early indicating a mean value of days 1, 2 and 3 post challenge. Comparisons were made with two-way ANOVA using Tukey’s multiple comparisons test. Asterisks represent significant comparisons (*, p<0.05; **, p<0.01;****, p<0.0001).

**Figure S4:**
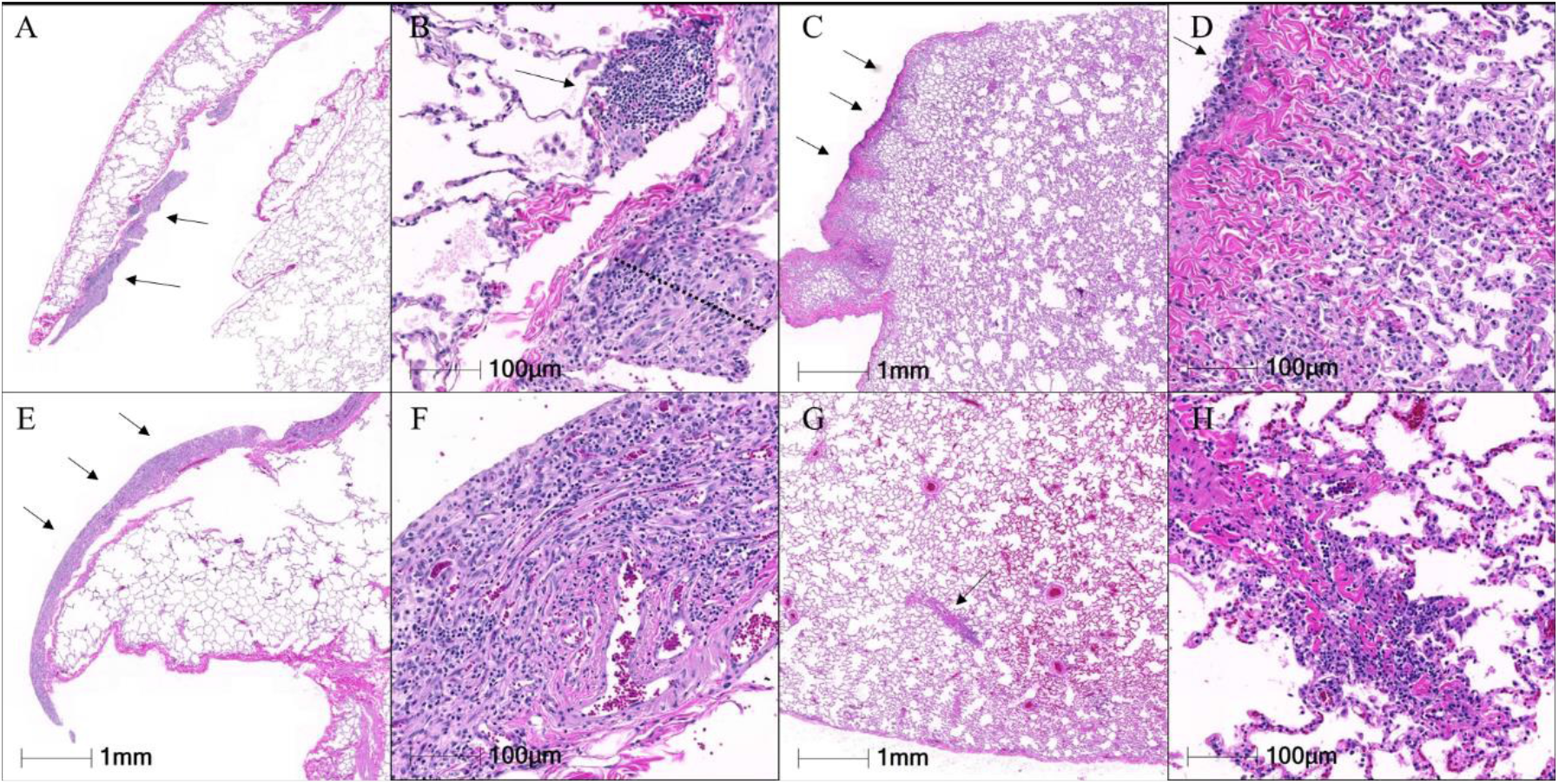
Representative histopathology. A,B: Aerosol RM, right middle lobe. A) The pleura is segmentally thickened (pleuritis, arrows). B) Regions of pleuritis are characterized by fibrosis (dotted line) with infiltration by mononuclear cells. Aggregates of similar inflammatory cells are present subpleurally (arrow). C,D: Aerosol AGM, left anterior lobe. C) The pleura is segmentally thickened (arrows). D) The pleura is lined by hypertrophic mesothelial cells (arrow) and there is infiltration of the subpleural parenchyma by histiocytes. E,F: IT/IN RM, left lower lobe. E) The pleura is segmentally thickened (pleuritis, arrows). F) The pleura is thickened by fibrosis and infiltrated by mononuclear cells, predominately lymphocytes. G,H: IT/IN AGM, right lower lobe. G) There is mild congestion and rare perivascular inflammation (arrow). H) Perivascular inflammation is characterized by infiltration of the tunica adventitia by mononuclear cells.

**Figure S5:**
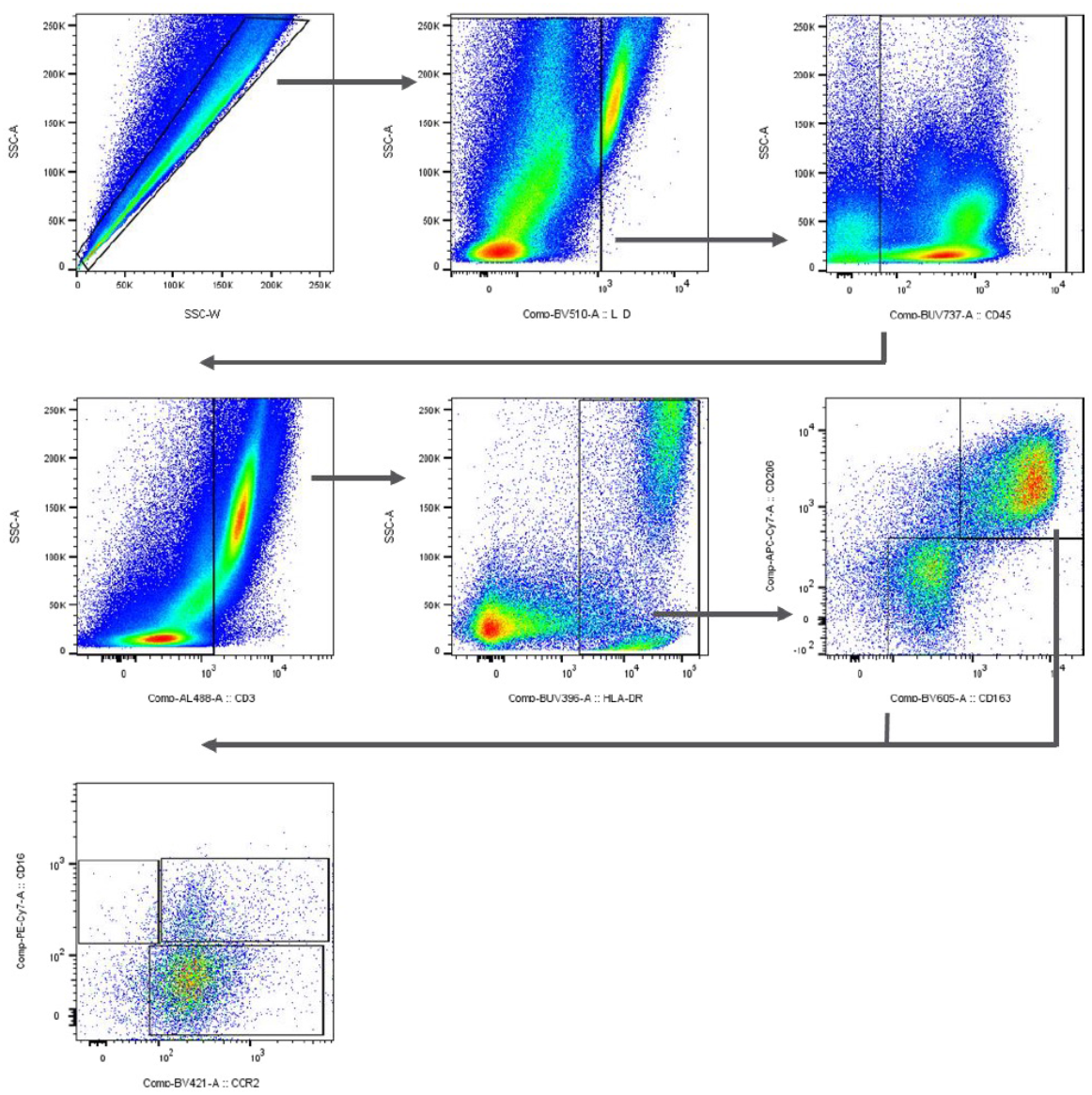
BAL flow cytometry gating strategy. Representative gating strategy to classify alveolar (CD163+CD206+), interstitial (CD163+CD206-), monocyte-derived (CD163+CD206+CD16+CCR2+), and resident alveolar (CD163+CD206+CD16-) macrophages in BAL.

**Figure S6:**
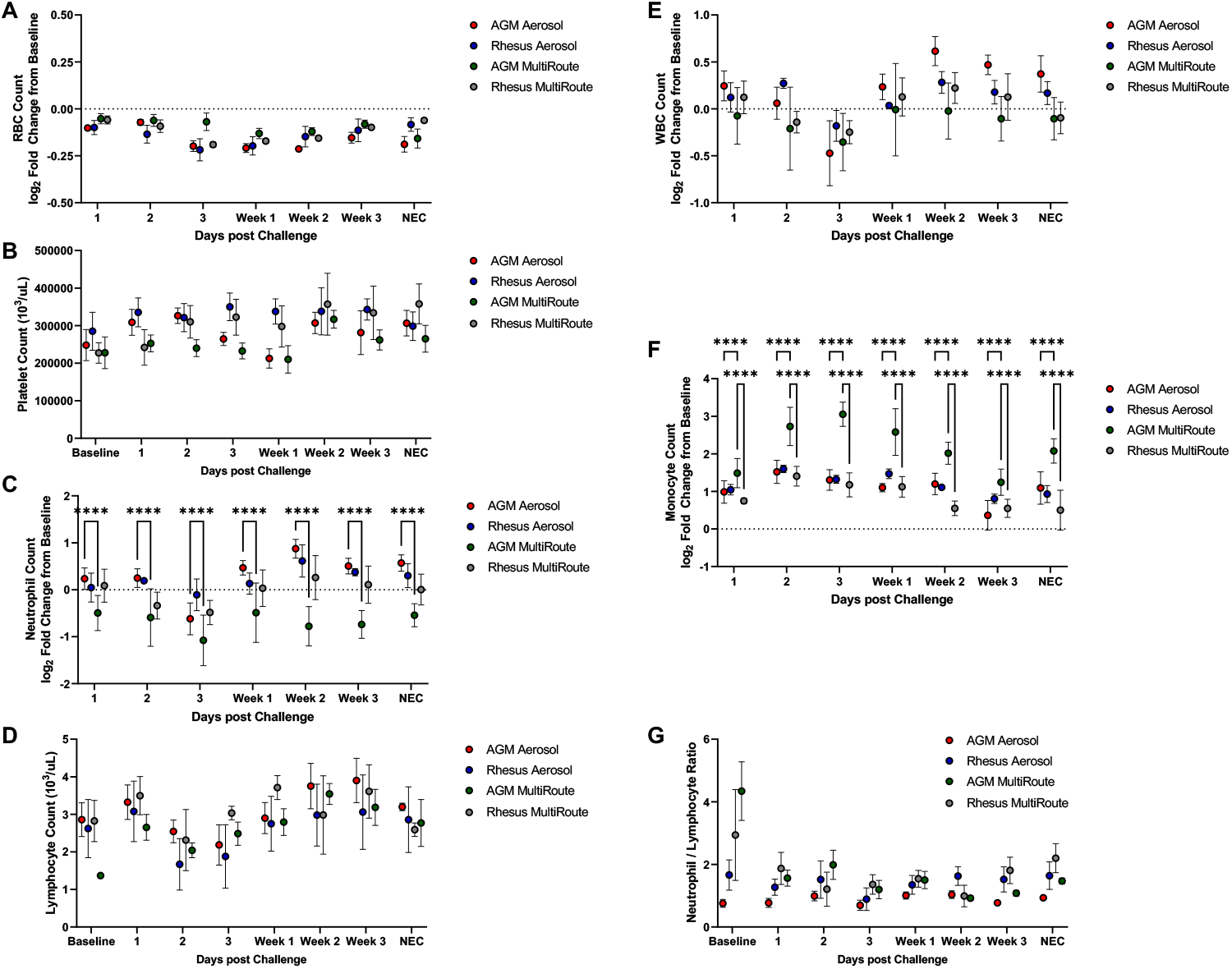
Hematology-Based Parameters of SARS-CoV-2 Challenge. Complete blood counts were performed at indicated times and were compared for counts of RBCs, platelets, neutrophils, lymphocytes, WBCs and monocytes (A, B, C, D, E, and F, respectively), as well as neutrophil/lymphocytes ratio (G). Comparisons were made via two-way ANOVA with Tukey’s multiple comparisons test. Asterisks represent significant comparisons (****, p<0.0001).

**Figure S7:**
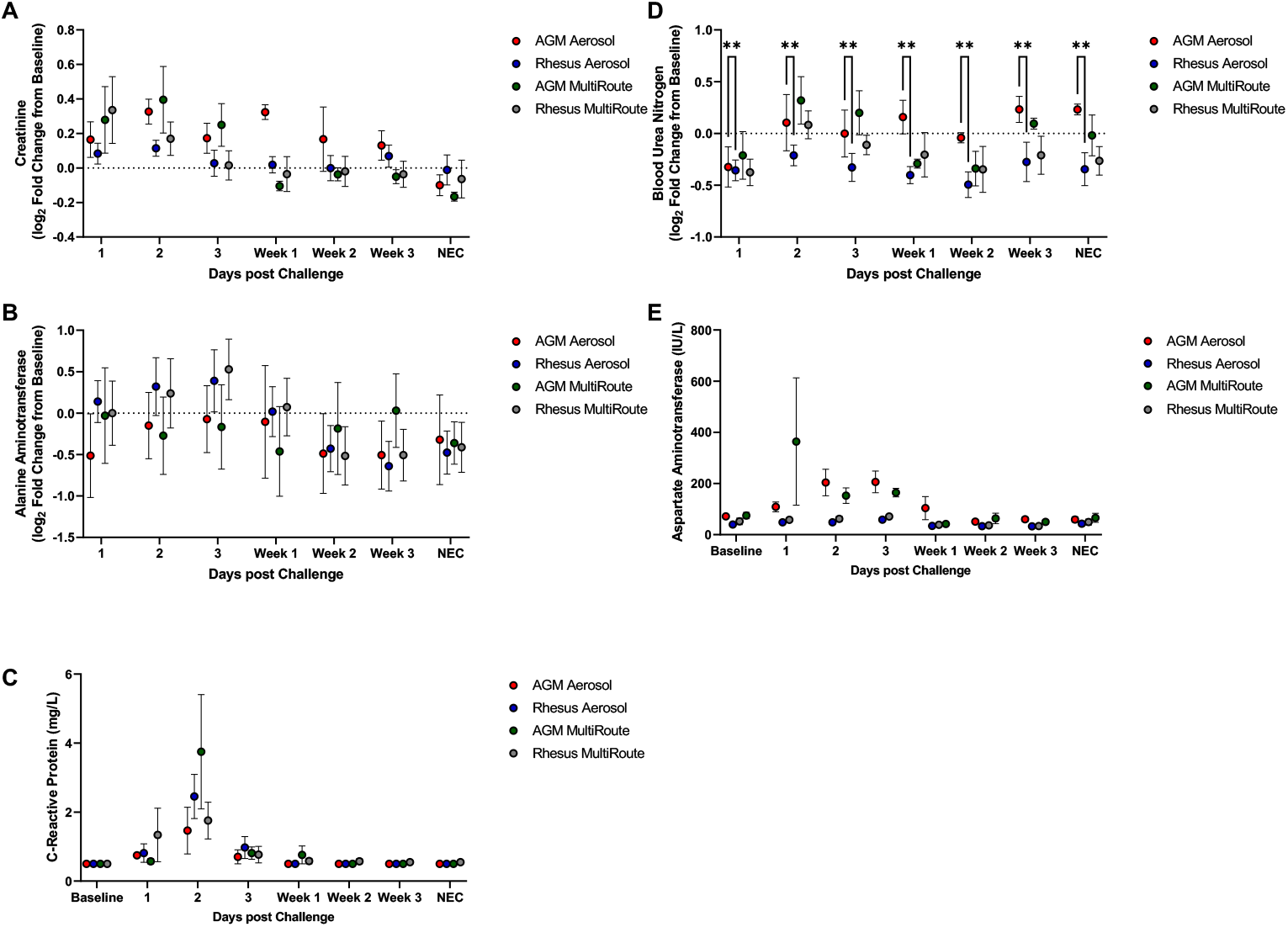
Clinical Chemistry-Based Parameters of SARS-CoV-2 Challenge. Clinical chemistries were performed at the indicated times post challenge. Comparisons between each group were made for log_2_ fold change from baseline of creatinine (A), ALT (B), BUN (D) and concentrations of CRP (C) and AST (E). Comparisons were made via two-way ANOVA with Tukey’s multiple comparisons test. Asterisks represent significant comparisons (**, p<0.01).

**Table S1:** Study Animal Characteristics.

| Animal | Species | Age (years) | Sex | Weight (kg) | Dose (TCID<sub>50</sub>) | Exposure Route |
| --- | --- | --- | --- | --- | --- | --- |
| IR12 | Macaca mulatta | 10 | Male | 9.5 | 1.9 x 10<sup>3</sup> | Aerosol |
| IJ01 | Macaca mulatta | 11 | Male | 10.4 | 2.2 x 10<sup>3</sup> | Aerosol |
| KN90 | Macaca mulatta | 7 | Male | 11.1 | 1.6 x 10<sup>4</sup> | Aerosol |
| JG28 | Macaca mulatta | 10 | Male | 12.1 | 7.5 x 10<sup>4</sup> | Aerosol |
| NB86 | Chlorocebus aethiops | 7 | Male | 7.1 | 7.1 x 10<sup>3</sup> | Aerosol |
| NB78 | Chlorocebus aethiops | 7 | Male | 6.3 | 4.4 x 10<sup>3</sup> | Aerosol |
| NB76 | Chlorocebus aethiops | 7 | Male | 6.6 | 6.0 x 10<sup>3</sup> | Aerosol |
| NC06 | Chlorocebus aethiops | 7 | Male | 6.0 | 5.0 x 10<sup>3</sup> | Aerosol |
| JK41 | Macaca mulatta | 9 | Male | 11.5 | 2.0 x 10<sup>6</sup> | Multi-route |
| KH77 | Macaca mulatta | 7 | Male | 13.0 | 2.0 x 10<sup>6</sup> | Multi-route |
| KT88 | Macaca mulatta | 7 | Male | 13.3 | 2.0 x 10<sup>6</sup> | Multi-route |
| LP05 | Macaca mulatta | 4 | Male | 7.0 | 2.0 x 10<sup>6</sup> | Multi-route |
| NB77 | Chlorocebus aethiops | 7 | Male | 5.4 | 2.0 x 10<sup>6</sup> | Multi-route |
| NC04 | Chlorocebus aethiops | 7 | Male | 5.9 | 2.0 x 10<sup>6</sup> | Multi-route |
| NC07 | Chlorocebus aethiops | 7 | Male | 6.4 | 2.0 x 10<sup>6</sup> | Multi-route |
| NB81 | Chlorocebus aethiops | 7 | Male | 6.3 | 2.0 x 10<sup>6</sup> | Multi-route |

**Table S2:**
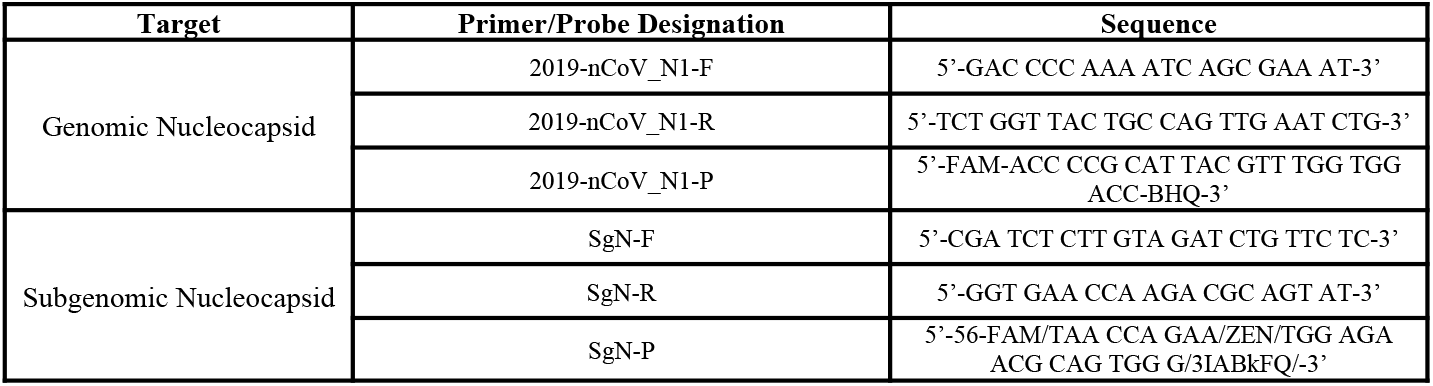
RT-qPCR Primers and Probes.

**Table S3:** Antibodies used for fluorescent immunohistochemistry.

| Target | Manufacturer | Catalog # | Use | Dilution |
| --- | --- | --- | --- | --- |
| Cytokeratin V | Abcam | ab17130 | Primary | 1:50 |
| p63 | GeneTex | GTX102425 | Primary | 1:100 |
| αSMA | Dako | M0851 | Primary | 1:100 |
| CD163 | Leica | NCL-L_CD163 | Primary | 1:50 |
| CD206 | R&D Systems | AF2535 | Primary | 1:50 |
| Collagen I | Abcam | ab34710 | Primary | 1:100 |
| Alexa 568 | Invitrogen | A-11036 | Secondary | 1:1000 |
| Alexa 568 | Invitrogen | A21134 | Secondary | 1:1000 |
| Alexa 568 | Molecular Probes | A11057 | Secondary | 1:1000 |
| Alexa 488 | Invitrogen | A-11034 | Secondary | 1:1000 |
| Alexa 488 | Invitrogen | A21202 | Secondary | 1:1000 |
| Alexa 488 | Invitrogen | A21121 | Secondary | 1:1000 |

**Table S4:**
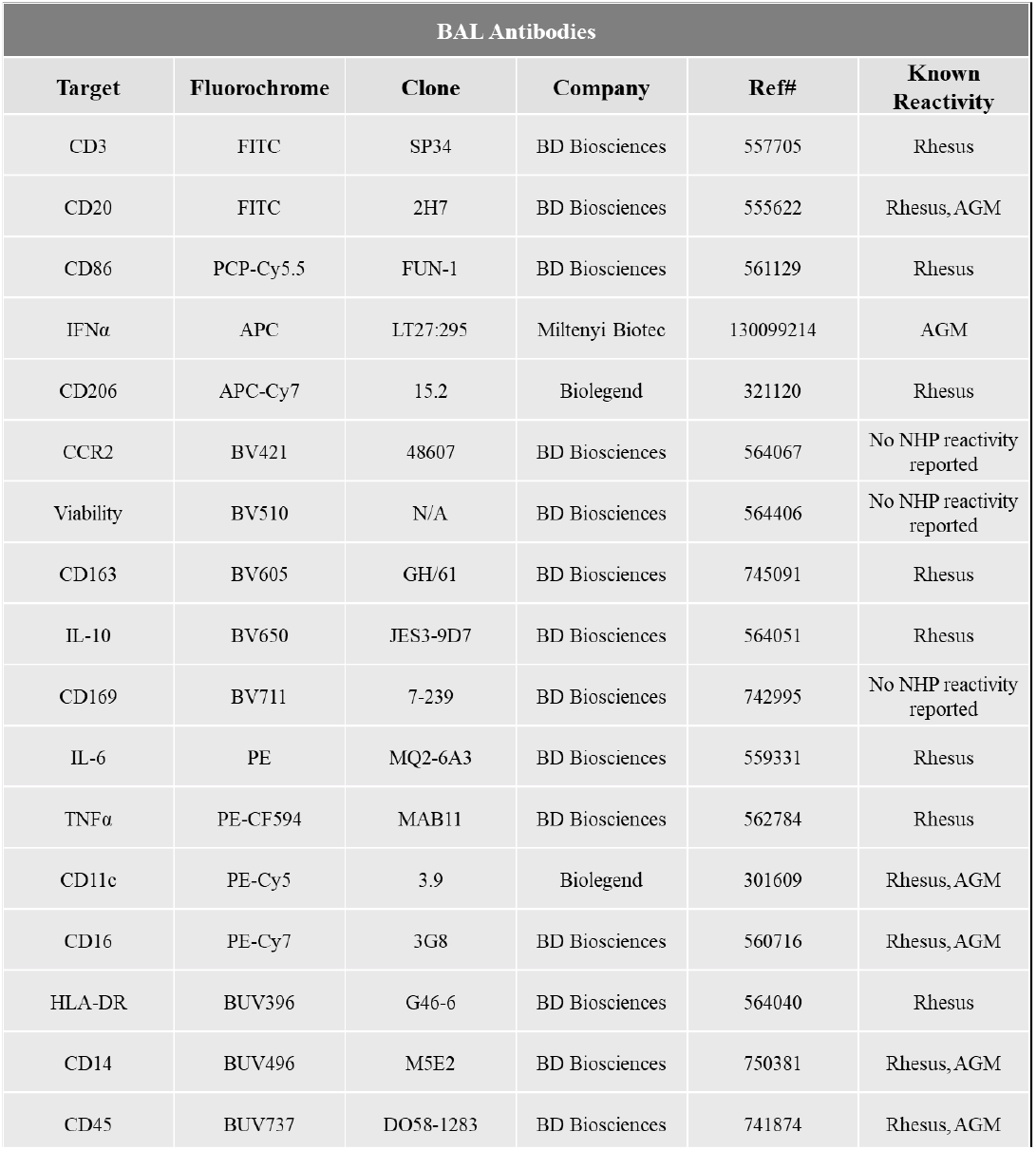
Antibodies used for flow cytometry analysis.

| BAL Antibodies |  |  |  |  |  |
| --- | --- | --- | --- | --- | --- |
| Target | Fluorochrome | Clone | Company | Ref# | Known Reactivity |
| CD3 | FITC | SP34 | BD Biosciences | 557705 | Rhesus |
| CD20 | FITC | 2H7 | BD Biosciences | 555622 | Rhesus, AGM |
| CD86 | PCP-Cy5.5 | FUN-1 | BD Biosciences | 561129 | Rhesus |
| IFNα | APC | LT27:295 | Miltenyi Biotec | 130099214 | AGM |
| CD206 | APC-Cy7 | 15.2 | Biolegend | 321120 | Rhesus |
| CCR2 | BV421 | 48607 | BD Biosciences | 564067 | No NHP reactivity reported |
| Viability | BV510 | N/A | BD Biosciences | 564406 | No NHP reactivity reported |
| CD163 | BV605 | GH/61 | BD Biosciences | 745091 | Rhesus |
| IL-10 | BV650 | JES3-9D7 | BD Biosciences | 564051 | Rhesus |
| CD169 | BV711 | 7-239 | BD Biosciences | 742995 | No NHP reactivity reported |
| IL-6 | PE | MQ2-6A3 | BD Biosciences | 559331 | Rhesus |
| TNFα | PE-CF594 | MAB11 | BD Biosciences | 562784 | Rhesus |
| CD11c | PE-Cy5 | 3.9 | Biolegend | 301609 | Rhesus, AGM |
| CD16 | PE-Cy7 | 3G8 | BD Biosciences | 560716 | Rhesus, AGM |
| HLA-DR | BUV396 | G46-6 | BD Biosciences | 564040 | Rhesus |
| CD14 | BUV496 | M5E2 | BD Biosciences | 750381 | Rhesus, AGM |
| CD45 | BUV737 | DO58-1283 | BD Biosciences | 741874 | Rhesus, AGM |

